# Antigen-agnostic identification of poxvirus broadly neutralizing antibodies targeting OPG153

**DOI:** 10.1101/2025.08.15.670377

**Authors:** Ida Paciello, Ling Zhou, Emily J. Rundlet, Giulia Realini, Federica Perrone, Giulio Pierleoni, Marco Troisi, Jeanne Postal, Florence Guivel-Benhassine, Françoise Porrot, Connor M. Mullins, Davide Moschese, Maria Vittoria Cossu, Federico Sabaini, Silvia Accordini, Natasha Gianesini, Rebeca Passarelli Mantovani, Francesca Panza, Massimiliano Fabbiani, Mario Tumbarello, Spinello Antinori, Concetta Castilletti, Francesca Montagnani, Olivier Schwartz, Rino Rappuoli, Jason S. McLellan, Emanuele Andreano

**Author notes:** Corresponding authors (R.R.); (J.S.M.); (E.A.). These authors contributed equally to this work (Ida Paciello, Ling Zhou, Emily J. Rundlet, Giulia Realini, Federica Perrone).

## Abstract

Recurrent mpox outbreaks caused by the monkeypox virus (MPXV) have prompted the World Health Organization to declare a Public Health Emergency of International Concern and have stimulated the development of medical interventions. Here, antigen-agnostic isolation of neutralizing monoclonal antibodies from convalescent or vaccinated people, AlphaFold 3-based predictive modeling, and cryo-electron microscopy were synergistically combined to identify the protein encoded by orthopoxviral gene (OPG) 153 (MPXV A28) as a target of broadly neutralizing antibodies. OPG153-targeting antibodies neutralized MPXV clade Ib, IIb, and vaccinia virus (VACV), and cross-reacted with OPG153 orthologs from cowpox and variola viruses. Immunization with MPXV OPG153 elicited a potent neutralizing antibody response against MPXV and VACV, substantiating OPG153 as a promising vaccine antigen and a potent target for preventive and therapeutic antibodies.

## Introduction

For more than five decades, monkeypox virus (MPXV) was considered a geographically restricted zoonotic pathogen, primarily affecting regions in Central (Clade I) and West (Clade II) Africa (*1*). However, beginning in 2022, MPXV exhibited unprecedented global human-to-human spread, predominated by the circulation of MPXV Clade IIb (MPXV-IIb) (*2, 3*). A subsequent outbreak in 2024, distinguished by the emergence of MPXV Clade Ib (MPXV-Ib), has further underscored the threat posed by this virus and prompted the World Health Organization (WHO) to declare a Public Health Emergency of International Concern (*2, 4*). From January 2022 through March 2025, the WHO reported 137,892 mpox cases globally and 317 deaths in 132 countries (*5*). As of April 2025, there have been over 30,000 estimated cases due to MPXV-Ib globally (*6*). The modified vaccinia Ankara-Bavarian Nordic (MVA-BN) vaccine—approved for emergency use during these outbreaks—demonstrated high real-world effectiveness (*7–12*). In smallpox vaccine–naïve individuals, however, a two-dose MVA-BN regimen induced modest and short-lived neutralizing antibody responses against MPXV (*10, 13, 14*). While the vaccine is generally well-tolerated, rare but severe adverse effects have been reported in immunocompromised individuals (*15*), warranting further clinical monitoring. Similarly, tecovirimat, the only approved antiviral for treating MPXV infection, did not reduce the duration of pox lesions among children and adults with MPXV-I infection in clinical trials (*16*). While current medical countermeasures have been instrumental in controlling recent mpox outbreaks, the development of next-generation vaccines and therapeutics with enhanced immunogenicity, durability, and clinical efficacy remains an unmet priority for global health.

MPXV is a large, double-stranded DNA virus within the *Orthopoxvirus* genus of the *Poxviridae* family and is genetically related to variola (VARV), vaccinia (VACV), and cowpox (CPXV) viruses (*1*). These are among the most complex human viruses known, encoding more than 200 gene products (*17*). Orthopoxviruses exist in two antigenically distinct virion forms: the intracellular mature virion (MV) and the extracellular enveloped virion (EV) (*2*). Prior studies identified several orthopoxvirus proteins targeted by neutralizing monoclonal antibodies (nAbs), including MV proteins encoded by orthopoxvirus genes (OPG) 95 (MPXV M1; VACV L1) (*18–20*), 108 (MPXV and VACV H3) (*20–22*), 120 (MPXV E8; VACV D8) (*20, 23*), 154 (MPXV A29; VACV A27) (*20, 23–26*), 155 (MPXV A30; VACV A28) (*27*), and EV proteins OPG161 (MPXV A35; VACV A33) (*20, 22*) and OPG190 (MPXV B6; VACV B5) (*19, 20, 23, 28, 29*) (reviewed in (*30, 31*); OPG nomenclature established in (*32*)). Antibodies targeting these proteins neutralize the corresponding virion forms (*20, 28, 33*). These observations have guided the design of new vaccines, which have demonstrated that immunization with a combination of antigens from both the MV and EV forms provides enhanced protection compared to monovalent formulations (*34, 35*). Despite ongoing efforts, MPXV continues to spread globally, underscoring the urgent need for novel immunotherapeutics and vaccine candidates. To address this, we combined Reverse Vaccinology 2.0 (*36*), AlphaFold 3-based predictive modeling (*37*), and electron microscopy to isolate and characterize nAbs from MPXV-infected or MVA-BN-vaccinated individuals, identify their antigenic target, and design an effective monovalent vaccine candidate.

## Results

Plasma and peripheral blood mononuclear cells (PBMCs) were collected from three individuals who recovered from MPXV-IIb infection during the 2022 outbreak and three healthy donors with a history of vaccination for VARV who received a booster dose of MVA-BN vaccine. The average age of the infected and vaccinated groups was 47 and 44 years, respectively, with sample collection occurring at an average of 100 days post-infection and 117 days post-vaccination (**fig. S1, A** and **B**). Plasma reactivity against UV-inactivated MPXV-IIb, VACV, and CPXV was assessed by ELISA, revealing cross-reactive binding in all donors (**fig. S1C**, *left*). Neutralization of live MPXV-IIb was quantified using a cytopathic effect-based microneutralization assay (CPE-MN) (*38, 39*), while live VACV neutralization was quantified using a virus-mediated U2OS cell-cell fusion assay (*40, 41*). Sera from the convalescent individuals exhibited neutralizing activity against both viruses, with titers ranging from 1:10 to 1:80 for MPXV-IIb and 1:160 to 1:320 for VACV (**fig. S1C**, *middle*). In contrast, two vaccinees displayed low VACV-neutralizing antibody titers (1:10 to 1:20), and none of the vaccinees showed cross-neutralization against MPXV-IIb, consistent with prior findings (*42*). Given the importance of Fc-mediated effector functions in antiviral immunity (*43*), we further assessed antibody-dependent cellular phagocytosis (ADCP) and antibody-dependent cellular cytotoxicity (ADCC) against MPXV-IIb. Infection elicited both ADCP and ADCC activity, whereas vaccinated donors showed limited Fc effector responses, with only one donor exhibiting detectable ADCC activity (**fig. S1C**, *right*).

We implemented an antigen-agnostic single-cell sorting strategy focused on CD19^+^CD27^+^IgD^−^IgM^−^ class-switched memory B cells (MBCs) to isolate MPXV-neutralizing antibodies (**Fig. 1A**). From these cohorts, a total of 69,300 MBCs were sorted, with frequencies of class-switched IgM^-^ cells ranging between 88.6% and 98.3% (**Fig. 1B**). Sorted cells were cultured over a layer of 3T3-CD40L feeder cells in the presence of IL-2 and IL-21 stimuli for 2 weeks to release antibodies into the supernatant (*38, 39, 44*). The supernatants from all cultured wells—containing immunoglobulin G (IgG) or A (IgA)—were screened via CPE-MN for neutralization of MPXV-IIb, which contained both MV and EV forms. This screen identified 43 nAbs (0.06%; **Fig. 1C**). From this panel of antibodies, we recovered 12 paired heavy chain and light chain variable regions, which were then expressed as recombinant human IgG1 for further characterization (**table S1**). Analysis showed broad diversity among immunoglobulin heavy chain variable (IGHV) and joining (IGHJ) gene usage, with no clonal lineages. In contrast, the light chains were skewed toward the IGKV3-20 gene, despite pairing with different J-genes. Notably, some heavy chains and most light chains were heavily mutated, with V gene mutation frequencies above 20%.

**Figure 1.**
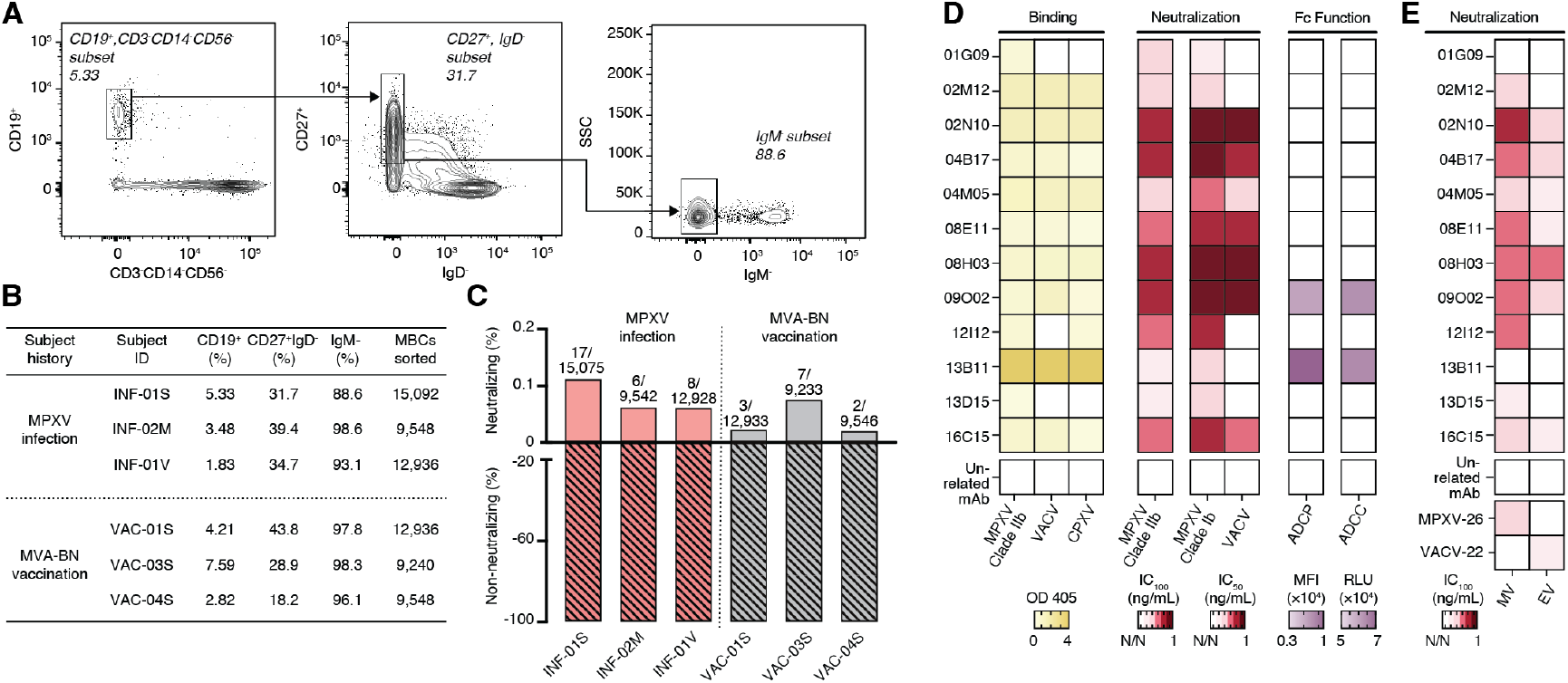
Antigen-agnostic single-cell sorting identified 12 MPXV nAbs. (**A**) Single-cell sorting strategy to isolate MPXV-neutralizing antibodies from class-switched memory B cells. From left to right: CD19^+^CD3^-^CD14^-^CD56^-^, CD27^+^IgD^-^, and IgM^-^ cells. (**B**) Frequencies of the B cell populations and the number of MBCs sorted from each donor enrolled in this study: three MPXV-infected donors and three MVA-BN vaccinees. (**C**) Proportion of neutralizing (solid) and non-neutralizing (diagonal line pattern) antibodies from MPXV-infected donors (pink) and MVA-BN vaccinees (gray). The number of neutralizing and non-neutralizing mAbs tested per individual is indicated. (**D**) Heatmaps showing: (*left*) binding activity to UV-inactivated MPXV, CPXV, and VACV; (*middle*) the neutralization potency against MPXV-IIb (IC_100_), MPXV-Ib (IC_50_), and VACV (IC_50_); and (*right*) the Fc-mediated effector functions (ADCP and ADCC) against UV-inactivated MPXV. (**E**) Heatmap showing neutralization potency (IC_100_) against MV and EV forms of MPXV-IIb. MPXV-26 (anti-OPG95; MPXV-M1; MV), VACV-22 (anti-OPG161; MPXV A35; EV) (*20*), and an unrelated mAb were used as controls.

The 12 recombinantly expressed antibodies were assessed by ELISA for binding to UV-inactivated MPXV-IIb, VACV, and CPXV. All antibodies bound MPXV-IIb; of these, one antibody (8.3%; 12I12) bound only to MPXV and CPXV, while nine (75%; 02M12, 02N10, 04B17, 04M05, 08E11, 08H03, 09O02, 13B11, and 16C15) showed cross-reactivity to VACV and CPXV (**Fig. 1D**, *left*). Neutralizing potency against MPXV-IIb was determined using the CPE-MN assay, with 100% inhibitory concentration (IC_100_) values ranging from 31.3 ng/mL to >2,000 ng/mL (**Fig. 1D**, *middle*, and **table S2**). Breadth of neutralization was further evaluated against MPXV-Ib and VACV. All antibodies neutralized MPXV-Ib, with half-maximal inhibitory concentration (IC_50_) values spanning 0.7 ng/mL to >23,000 ng/mL. Seven antibodies (58.3%; 02N10, 04B17, 04M05, 08E11, 08H03, 09O02, and 16C15) exhibited VACV cross-neutralization, with IC_50_ values between 5.7 and 1,058 ng/mL. To explore Fc-effector function potential, we assessed ADCP and ADCC activity. Among the 12 antibodies, only 09O02 and 13B11 elicited both ADCP and ADCC responses (**Fig. 1D**, *right*). Next, we investigated the neutralization of MPXV virion forms. To control for cross-contamination of viral forms, we confirmed that each form was neutralized only by its respective control nAb (VACV-22 for EV and MPXV-26 for MV). Three antibodies (25.0%; 02M12, 12I12, and 13D15) showed MV-specific neutralization while, unexpectedly, seven antibodies (58.3%; 02N10, 04B17, 04M05, 08E11, 08H03, 09O02, and 16C15) neutralized both MV and EV (**Fig. 1E** and **table S2**). Two antibodies (16.7%; 01G09 and 13B11) lacked detectable activity against either form, which may reflect their lower neutralization potency.

Collectively, these results define diverse functional profiles and crosspoxvirus reactivity within the antibody panel.

MPXV has >35 annotated surface-exposed proteins, most of which have unknown structures or functions. Therefore, to narrow the pool of potential targets of the nAbs isolated from donor samples, we used AlphaFold 3 (*37*) to screen for predicted pairwise interactions with MPXV full-length envelope proteins. High-confidence interactions between OPG153 (MPXV A28; VACV A26) and nAbs 08E11 and 12I12 were predicted, with interface predicted template modelling (iPTM) scores ≥0.85 (**Fig. 2A**).

**Figure 2.**
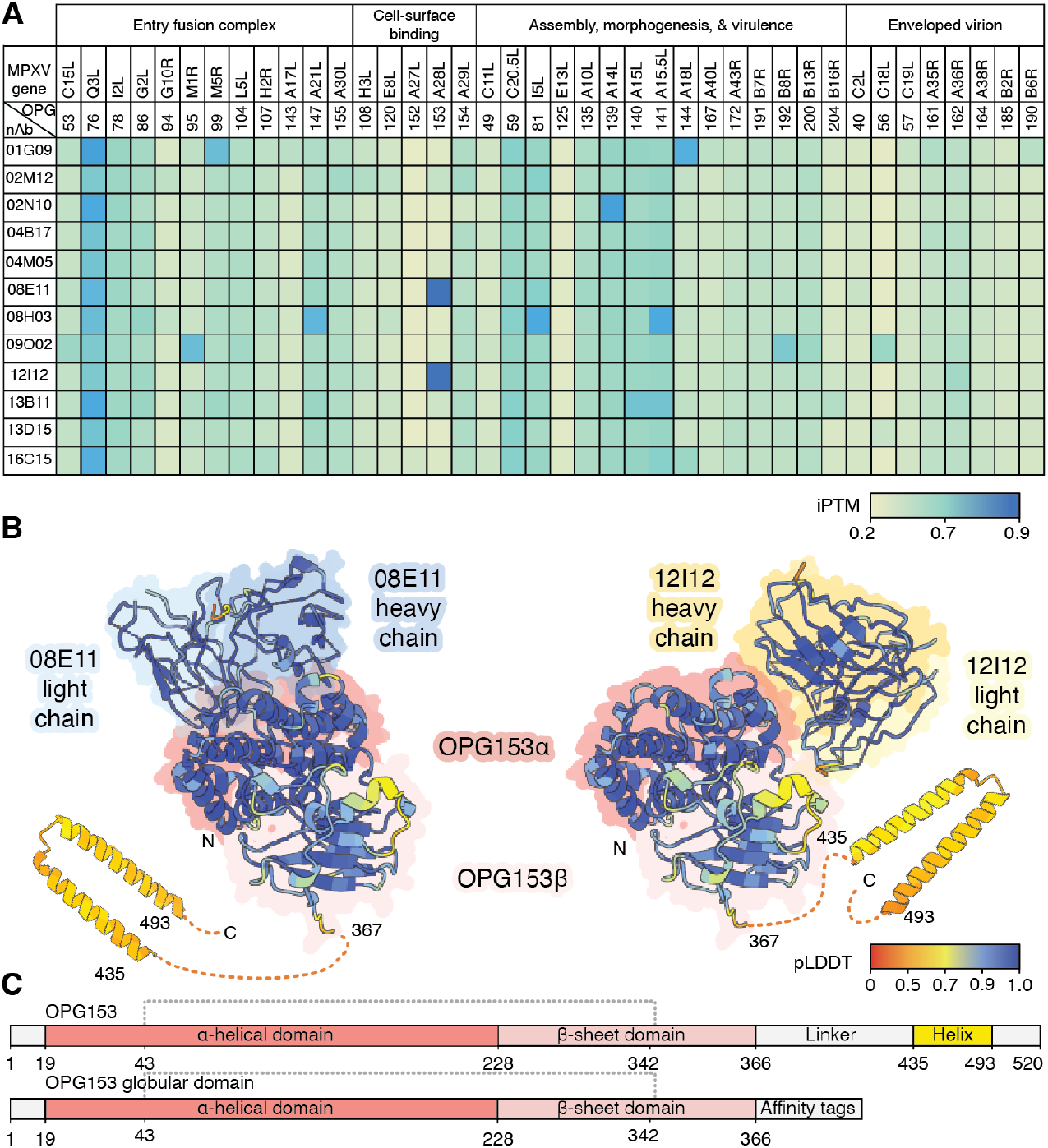
AlphaFold 3-based screening identifies OPG153 as a target of MPXV-neutralizing antibodies. (**A**) Heatmap of AlphaFold 3 interface predicted template modeling (ipTM) scores between annotated MPXV surface antigens and nAbs isolated from donor samples (*37*). (**B**) Predicted structural models of MPXV OPG153 (A28L) in complex with nAbs 08E11 (blue, *left*) and 12I12 (yellow, *right*), colored by predicted local distance difference test (pLDDT) scores. Disordered regions of OPG153 (residues 367–435 and 493–520) are omitted for clarity. (C) Schematic representation of full-length OPG153 (*top*) and the truncated expression construct encoding the N-terminal globular domain (*bottom*). Dotted line indicates a disulfide bond.

08E11 and 12I12 were predicted to interact with the N-terminal globular domain of OPG153 (**Fig. 2, B** and **C**), recognizing non-overlapping epitopes (**fig. S2**). OPG153 plays multifaceted roles in orthopoxvirus biology, with evidence suggesting it mediates (i) host interaction (*45*), (ii) A-type inclusion body formation (*46, 47*), (iii) EV generation (*48, 49*), and (iv) cell entry (*50–52*). Its association with the viral membrane is mediated by a C-terminal, disulfide-bonded, coiled-coil interaction with OPG154 (MPXV A29; VACV A27) (*48, 53*). OPG153 has been reported to associate exclusively with the MV form of VACV (*49*), highlighting the need for further investigations into differences in surface antigen presentation between MV and EV forms.

Based on the AlphaFold 3 structural predictions identifying MPXV OPG153 as a potential target of nAbs, we expressed and purified the globular domain (residues 1–366) with a C-terminal affinity tag (**Fig. 2C** and **fig. S3**). OPG153 does not possess a signal peptide, yet we purified this protein from the supernatant of transiently transfected FreeStyle 293-F cells (**fig. S3**). Surface plasmon resonance (SPR) experiments showed high-affinity binding for the antigen-binding fragments (Fabs) of 08E11 and 12I12 to MPXV OPG153 N-terminal domain, with *K*_D_s of 18.5 nM and 3.2 nM, respectively (**fig. S4**). These binding results validated the AlphaFold 3 hits, supporting that iPTM scores ≥0.85 can be used to identify antibody-antigen interactions and that OPG153 is a target of MPXV-neutralizing antibodies.

We next performed a flow cytometry-based assay to assess the binding of all 12 nAbs to OPG153. Notably, we found that in addition to 08E11 and 12I12, six more nAbs (02N10, 04B17, 16C15, 08H03, 09O02, and 02M12) bound to the OPG153 N-terminal domain (**Fig. 3A**, *left*). The AlphaFold 3-predicted interactions between OPG153 and these six additional antibodies were of low confidence (**Supplementary Data File 1**). Given the high sequence conservation of OPG153 among orthopoxviruses (>93% identity; **fig. S5**), we further evaluated cross-reactivity to the OPG153 orthologs from VACV, CPXV, and VARV. All MPXV OPG153-reactive nAbs bound the VACV and CPXV OPG153 orthologs, with the exception of 12I12, which did not interact with VACV OPG153 (**Fig. 3A**, *left*, and **fig. S4**). These data are consistent with the observed lack of VACV neutralization or OPG153-binding by 12I12 (**Fig. 1D** and **fig. S4**). Similarly, all MPXV OPG153-binding antibodies, except 08E11, interacted with OPG153 of VARV (**Fig. 3A**, *left*). Four antibodies (01G09, 04MO5, 13B11, and 13D15) did not bind any tested OPG153 orthologs and are presumed to target different MPXV antigens.

**Figure 3.**
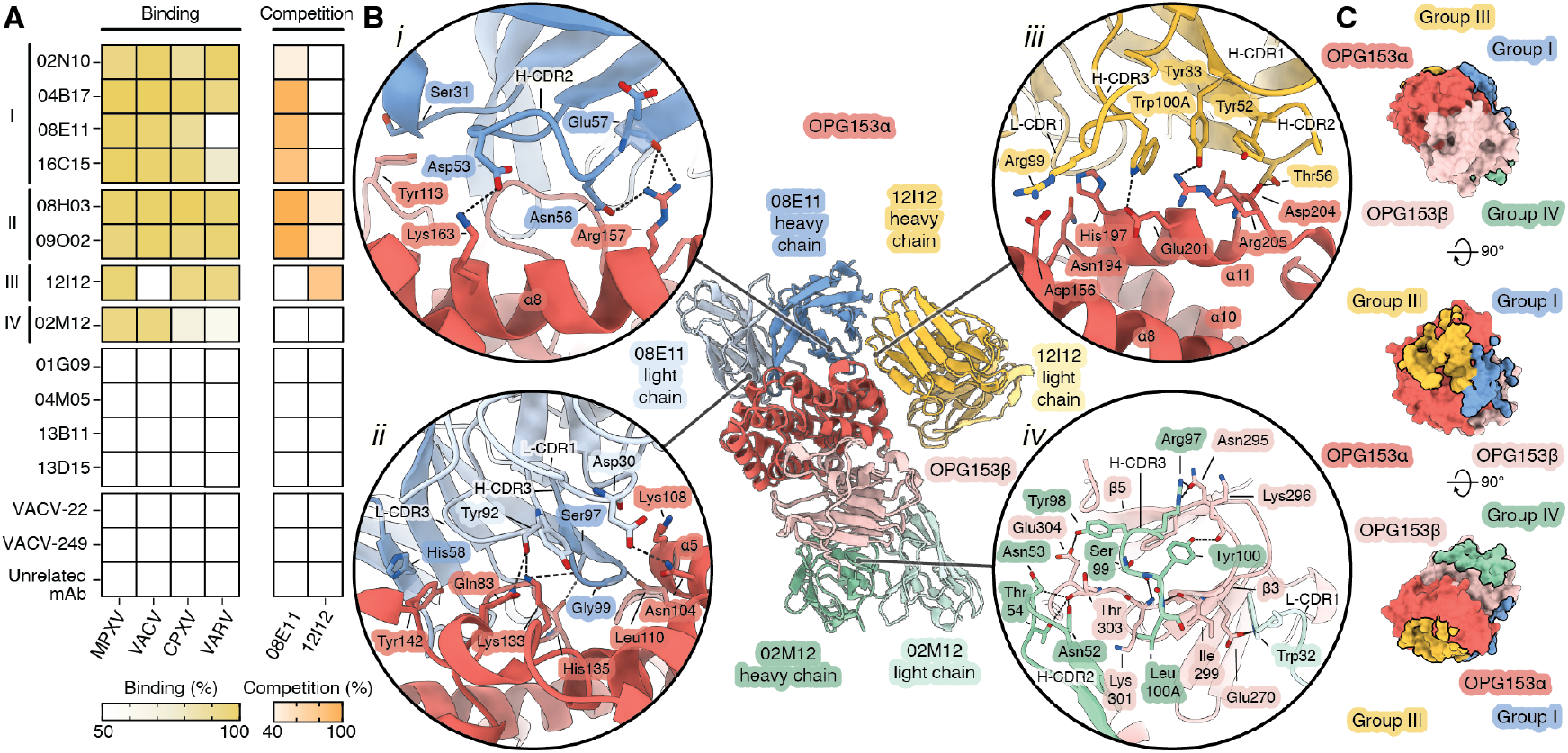
OPG153-specific nAbs recognize distinct antigenic sites. (**A**) Heatmaps showing the flow cytometry-based binding activity (*left*) and competition against 12I12 or 08E11 (*right*) for the 12 nAbs. Antigen binding was defined as occurring when >50% of the bead population was Alexa488^+^; competition was determined by a >40% reduction in fluorescent signal. Antibodies VACV-22 (anti-OPG161; MPXV A35), VACV-249 (anti-OPG120; MPXV-E8) (*20*), and an unrelated mAb were used as negative controls. (**B**) Cryo-EM structural models of MPXV OPG153 bound to Fabs from 08E11 (blue), 12I12 (yellow), and 02M12 (green). Insets display close-up views of molecular interactions, including hydrogen bonds and salt bridges (dotted black lines). Oxygen atoms are red and nitrogen atoms are blue. (**C**) Surface representation of OPG153 showing epitope footprints for 08E11 (group I), 12I12 (group III), and 02M12 (group IV).

To evaluate whether the OPG153-binding antibodies recognized overlapping or distinct epitopes, we conducted competition assays using labeled 08E11 and 12I12 as competitors, which we chose due to their different predicted binding epitopes and neutralization profiles. Based on binding-interference patterns, antibodies were classified into four competition groups (**Fig. 3A**). Group I antibodies (02N10, 04B17, 08E11, and 16C15) competed with 08E11 (**Fig. 3A**, *right*). These antibodies displayed broad binding to OPG153 orthologs and potent neutralization across MPXV clades and virion forms, as well as VACV (**Figs. 1, D** and **E**, and **3A**, *right*). Group II antibodies (08H03 and 09O02) competed with both 08E11 and 12I12 and shared the same binding and neutralization profiles as Group I antibodies. Group III antibody, 12I12, competed with itself but not with 08E11. Group IV antibody, 02M12, did not compete with 08E11 nor 12I12, suggesting it targets a distinct epitope on OPG153. SPR studies showed that 02M12 also binds with high-affinity to MPXV OPG153 (*K*_D_ = 30.7 nM; **fig. S4**). Despite its broad OPG153 reactivity, 02M12 exhibited lower neutralization breadth, with no activity against VACV or MPXV-IIb EV (**Fig. 1, D** and **E**), suggesting its epitope may be less neutralization-sensitive.

To investigate the interactions between the nAbs and MPXV OPG153, we conducted cryo-electron microscopy (cryo-EM) analyses using a 200 kV Glacios microscope. Guided by the AlphaFold 3-predictions and our binding studies, we assembled a ternary complex of OPG153 N-terminal domain with Fabs 08E11 and 12I12 and a binary complex of OPG153 with Fab 02M12. For the ternary complex, two datasets comprising 891 movies at 0° tilt and 1,147 movies at 30° tilt were acquired and merged for processing (**fig. S6** and **table S3**). A total of 186,643 particles were selected for the final 3D reconstruction, yielding a 2.8 Å resolution map. For the binary complex, 1,743 movies were collected, resulting in 132,110 final particles that yielded a 3.2 Å resolution map (**fig. S7** and **table S3**). The refined structures of the MPXV OPG153 N-terminal domain agree well with the AlphaFold 3 predictions (R.M.S.D. = 0.4 Å and 0.5 Å for 347 Cα atom pairs for the OPG153-08E11-12I12 and OPG153-02M12 structures, respectively; **fig. S8**) and with a 1.2 Å resolution crystal structure of the VACV OPG153 ortholog (R.M.S.D. = 0.5 Å for 347 Cα atom pairs; **fig. S9A**) (*54*). Neither EM map showed detectable N-acetylglucosamine map features at the three putative N-X-S/T glycosylation sequons (residues 194, 320, and 351; **fig. S9B**), consistent with the absence of a signal peptide in MPXV OPG153.

The cryo-EM map and model of the OPG153-08E11-12I12 complex reveal two adjacent but non-overlapping epitopes within the globular α-helical domain of MPXV OPG153 (**Fig. 3, B** and **C**, and **fig. S9C**, *top*) consistent with the AlphaFold 3 predictions (**fig. S8, A** and **B**). Fab 08E11 binding to OPG153 is mediated predominantly by its heavy chain, which buries 801 Å^2^ on the OPG153 surface. H-CDR2 makes the most contacts of the CDR loops, interacting mainly with α helix (α)7 and residues 156–167 within the α8 helix (**Fig. 3B**, *inset i*, and **figs. S5** and **S9D**) (*54*). The 08E11 light chain buries a more modest 315 Å^2^ of surface area on OPG153, with interactions primarily involving L-CDR1 and L-CDR3, contacting helices α4 and α5 and the linker between α6 and α7 (**Fig. 3B**, *inset ii*, and **figs. S5** and **S9D**). The 12I12 heavy chain buries a total surface area of 452 Å^2^ on the OPG153 globular α-helical domain (**Fig. 3B**, *inset iii*, and **figs. S5** and **S9E**). All three heavy chain CDR loops interact with OPG153, with H-CDR2 and H-CDR3 engaging OPG153 residues 197–210 within the α10 and α11 helices. Notably, the 12I12 H-CDR3 residue Arg99 (Kabat numbering) extends into the interface between all three proteins in the complex, establishing a salt bridge with OPG153 Asp156 in α8. The 12I12 light chain CDRs bury only 249 Å^2^ on OPG153, interacting with the N terminus of OPG153 α10 and the α13 helix.

02M12 buries a total of 876 Å^2^ of surface area on the β-sandwich domain of OPG153, distal to the 08E11 and 12I12 binding sites (**Fig. 3, B** and **C**, and **fig. S9C**, *bottom*). This interaction was not predicted by AlphaFold 3 (**fig. S8C**) and is predominantly mediated by 02M12 heavy-chain contacts with the loop of OPG153 between β-strands (β)5 and 6 (residues 295–305; **Fig. 3B**, *inset iv*, and **figs. S5** and **S9F**). Arg97 of H-CDR3 hydrogen bonds to Asn295 at the N terminus of the OPG153 loop, while adjacent H-CDR3 residue Tyr98 reaches in the opposite direction to interact with OPG153 Glu304. Residues 99–100A of H-CDR3 establish a series of antiparallel backbone interactions with OPG153 residues 299–303, while the sidechain of H-CDR3 Tyr100 also hydrogen bonds with the backbone of OPG153 Lys296. Light-chain contacts bury 277 Å^2^ of surface area on OPG153 and include Trp32— positioned C-terminal to L-CDR1—interacting with OPG153 Glu270 on β3. The reduced neutralization potency of 02M12 relative to 08E11 and 12I12 may indicate that the β-sandwich domain targeted by 02M12 is less neutralization-sensitive than the α-helical domain targeted by 08E11 and 12I12. This could imply that the α-helical domain may be more important for the function of OPG153, or that this domain is more accessible on the virion surface.

The identification of OPG153 as a major target of MPXV-neutralizing antibodies prompted us to evaluate the frequency of OPG153-specific MBCs in the infected and vaccinated individuals enrolled in this study. PBMCs were stained with Alexa Fluor 488-labeled recombinant MPXV OPG153 and analyzed by flow cytometry to detect CD19^+^CD27^+^IgD^−^IgM^−^OPG153^+^ MBCs. Antigen-specific MBCs were detected in all donor samples (**fig. S10**). However, MPXV-infected donors exhibited a 2.2-fold higher frequency of OPG153-specific MBCs than MVA-BN-vaccinated individuals, suggesting that natural infection elicits a more abundant response against this antigen or that the responses to MVA-BN antigens are lower overall.

To evaluate the immunogenicity of MPXV OPG153, 7–8-week-old BALB/c mice were immunized intramuscularly with 10 µg of MPXV-Ib OPG153 globular domain emulsified in 25 µL of MF59-like adjuvant (AddaVax; **Fig. 4A**). This group was compared to mice immunized with 50 µL of MVA-BN vaccine (infection units ranging from 0.5 to 3.95 × 10^7^) or 10 µg of severe acute respiratory syndrome coronavirus 2 (SARS-CoV-2) spike (S) protein (*55*) plus AddaVax. Each group consisted of 10 mice. Immunizations were administered on days 0 and 21, and serum samples were collected two weeks post-second dose (day 35). Neutralizing antibody titers were evaluated against MPXV-IIb containing both forms of the virus, MPXV-IIb MV, MPXV-IIb EV, and VACV (**Fig. 4, B** and **C**). Neutralization titers were reported as 100% inhibitory dilution (ID_100_) for MPXV and 50% inhibitory dilution (ID_50_) for VACV. As expected, sera from mice immunized with the SARS-CoV-2 S protein exhibited no neutralizing activity against MPXV-IIb or VACV. Conversely, OPG153 immunization elicited robust MPXV-IIb serum neutralization titers (median ID_100_ = 2,240), which were significantly higher than MVA-BN immunization (median ID_100_ = 280) (**Fig. 4B**, *left*). Compared to MVA-BN, immunization with OPG153 induced higher neutralizing titers against MV but lower titers against EV (**Fig. 4B**, *middle* and *right*). These data suggest that surface exposure and availability of OPG153 may differ between the two viral forms. Finally, immunization with OPG153 elicited significantly higher neutralization titers against VACV (median ID_50_ = 5,120) compared to MVA-BN immunization (median ID_50_ = 960), highlighting the potential of OPG153 to induce cross-poxvirus neutralizing antibodies (**Fig. 4C**).

**Figure 4.**
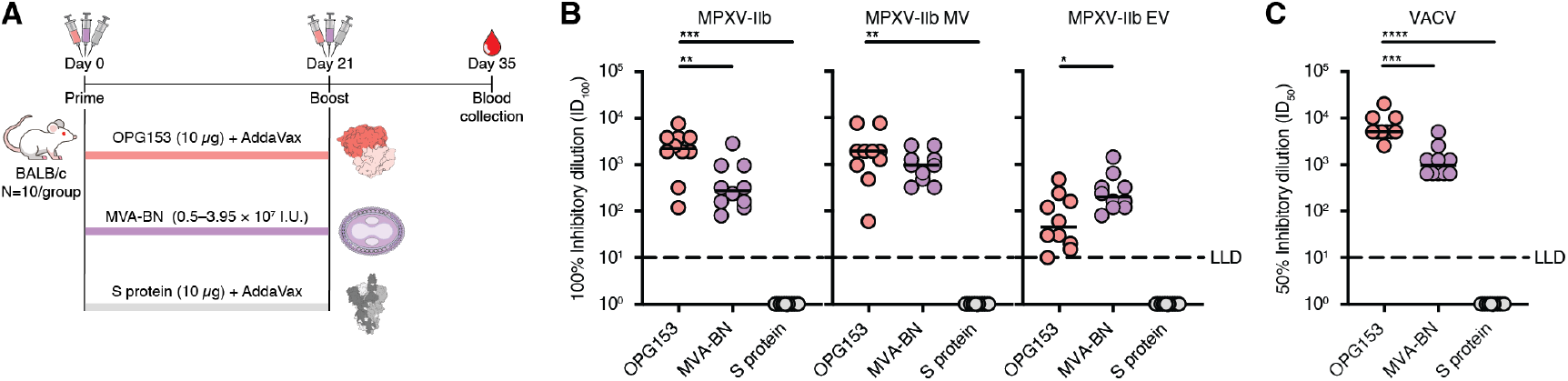
OPG153 immunization elicits cross-neutralizing antibody responses in BALB/c mice. (**A**) Schematic overview and timeline of the immunization study performed in BALB/c mice. (**B**) Neutralization titers elicited by immunization with OPG153, MVA-BN, or SARS-CoV-2 S protein, measured against MPXV-IIb virus (comprising both MV and EV forms, *left*), MPXV-IIb MV (*middle*), and MPXV-IIb EV (*right*). (**C**) Neutralization titers elicited against VACV. Statistical significance among groups was established using Tukey’s multiple comparisons test one-way ANOVA (95% confidence interval). Serum neutralization titers are shown as Log_10_-transformed values. Black lines indicate group medians; dotted lines represent the lower limit of detection (LLD).

## Discussion

The antigen-agnostic and AI-assisted approach described in this study led to the identification of OPG153 as a major target of orthopoxvirus broadly neutralizing antibodies. OPG153-targeting antibodies show high neutralization potency, bind at least three different regions of the OPG153 globular domain, and appear to neutralize both the MV and EV forms of the virus, although the dose required to neutralize EV was much higher than that needed to neutralize MV. This finding was unexpected as OPG153 has been reported to be associated exclusively with the MV form of VACV (*49*), necessitating further studies to address the biological role of OPG153 and its localization on MPXV virions. Nonetheless, our work suggests that OPG153 represents a promising target for preventive and therapeutic monoclonals and vaccines for MPXV and potentially other pathogenic orthopoxviruses.

Consistent with our findings, Tai et al. reported a systematic mRNA immunization screen of 34 MPXV surface proteins and found that full-length OPG153 was able to induce nAbs against MPXV, although neutralization was observed only against MV (*52*). They further demonstrated that an mRNA vaccine encoding OPG153 and 11 other MPXV antigens conferred protection against MPXV challenge in mice (*52*). In our study, immunization with the purified N-terminal domain of OPG153 as a monovalent subunit vaccine antigen elicited nAb titers comparable to those recently reported for multivalent mRNA vaccines (*7, 52, 56*), and significantly higher than those achieved following MVA-BN immunization (**Fig. 4**).

Collectively, this work supports OPG153 as a conserved and compelling target for next-generation orthopoxvirus vaccines. Monoclonal antibodies and single-antigen vaccines that protect across clades and morphologies represent promising orthopoxvirus therapeutics and may reduce manufacturing costs, thereby allowing for equitable access to low-and middle-income countries. Finally, the antibody- and antigen-discovery approach described here shows how AI-driven approaches can significantly accelerate the identification and development of unexploited targets for vaccines and therapeutics. Similar approaches can be used for emerging viral pathogens, antimicrobial-resistant bacteria, fungi, and parasites, leading to highly effective and affordable countermeasures for worldwide deployment.

## Acknowledgments

The authors acknowledge the Istituto Zooprofilattico Sperimentale della Puglia e della Basilicata (IZSPB) for sharing the CPXV Taunton strain, VisMederi Srl for the fruitful scientific discussions at the beginning of the project, and Farefarma Srl for the support with the *in vivo* experiments. The authors thank Drs. Axel Brilot and Evan Schwartz for their technical support at the Sauer Structural Biology Laboratory, The University of Texas at Austin. The authors thank Drs. Mahtab Beikzadeh and Kaci Erwin, and Jeong Ryeol Kim for FreeStyle 293-F cell culture support. The authors thank Ajit Ramamohan for support with cryo-EM data collection.

## Funding

The O.S. lab is funded by Institut Pasteur, Fondation pour la Recherche Médicale (FRM), ANRS, the Vaccine Research Institute (VRI) (ANR-10-LABX-77), Labex IBEID (ANR-10-LABX-62-IBEID), the HERA projects DURABLE (grant 101102733) and LEAPS. J.P. is supported by DURABLE. This work was funded in part by the Welch Foundation grant number F-0003-19620604 (J.S.M.). E.J.R. is a Fellow of The Jane Coffin Childs Fund for Medical Research.

## Author contributions

Study conception: R.R., J.S.M., and E.A.

Project coordination: R.R., J.S.M., and E.A.

Donor enrollment and blood collection: D.M., M.V.C, F.S., S.Ac., N.G., R.P.M., F.Pa., M.F., M.Tu.,

S.An., C.C. and F.M.

PBMC isolation: I.P.

Single cell sorting: I.P., G.R., F.Pe., and M.Tr.

MPXV-IIb neutralization screening: I.P., G.R., F.Pe., G.P., M.Tr., and E.A.

Antibody VH-VL recovery, cloning and expression: G.R. and F.Pe.

MPXV-IIb, MPXV-IIb EV, MPXV-IIb MV, MPXV-Ib, and VACV neutralization assays: I.P., G.R.,

F.Pe., J.P., F.G.B., and F.Po.

UV-inactivated MPXV-IIb, VACV, and CPXV ELISA: I.P.

Evaluation of antibody Fc function activities: I.P.

AlphaFold 3 screening: L.Z., E.J.R., and C.M.M.

FreeStyle 293-F cell culture and transient transfection: L.Z. and E.J.R.

Construct design, cloning, and protein purification: L.Z., E.J.R., G.R., and C.M.M. SPR: L.Z. and E.J.R.

Flow cytometry-based binding and competition assays: I.P.

Cryo-EM sample preparation, screening, and data collection: L.Z. and E.J.R.

Cryo-EM data processing and model building: L.Z., E.J.R., and J.S.M.

Manuscript writing and figure preparation: I.P., L.Z., E.J.R., G.R., F.Pe., R.R., J.S.M., and E.A.

All authors reviewed the final version of the figures and manuscript.

## Competing interests

R.R. holds shares in Novartis and GSK group of companies. I.P., G.R., F.Pe., G.P., M.Tr., R.R., and E.A. are inventors on Italian patent application no. 102024000030036 (Human monoclonal antibodies against Monkeypox and other Orthopoxviruses) filed on December 27, 2024. I.P., L.Z., E.J.R., G.R., F.Pe., R.R., J.S.M., and E.A. are inventors on US patent application no. UTSB.P1397US.P1 filed on May 9, 2025 (Monkeypox virus OPG153 globular domain vaccine).

## Data and materials availability

The structural models generated in this study have been deposited in the Protein Data Bank (PDB) (https://www.rcsb.org/). The corresponding Cryo-EM maps are available in the Electron Microscopy Data Bank (EMDB) (https://www.emdataresource.org/). All remaining data associated with this study are in the paper or supplementary materials.

## Supplementary Materials

Materials and Methods

Figs. S1 to S10

Tables S1 to S3

