## Supplemental Information for "Antigen-agnostic identification of poxvirus broadly neutralizing antibodies targeting OPG153"

**The PDF file includes:**
Materials and Methods
Figs. S1 to S10
Tables S1 to S3
References

#### Materials and Methods

Donor enrollment and human sample collection

Human samples from monkeypox virus (MPXV) convalescent donors and from individuals with MVA-vaccination were collected through a collaboration with the Azienda Ospedaliera Universitaria Senese, Siena (IT), the ASST FBF Sacco hospital, Milan (IT) and IRCCS Ospedale Sacro Cuore Don Calabria, Verona (IT). All the subjects enrolled gave their written consent. The study was approved by local ethics committees (Protocol numbers 22717 and 42936) and conducted according to good clinical practice in accordance with the Declaration of Helsinki (European Council 2001, US Code of Federal Regulations, ICH 1997). This study was unblind and not randomized. Three MPXV convalescent donors and three MVA-BN vaccinees were enrolled, then peripheral blood mononuclear cells (PBMCs) and plasma samples were collected. Plasma samples were heat-inactivated for 30 min at 56 °C for downstream assays.

Animal husbandry and welfare, and animal model ethics statement

Animal husbandry and experimental procedures were ethically reviewed and carried out in accordance with European Directive 2010/63/EU and Italian Decree 26/2014. The Facility is authorized for the use of animals for scientific procedures by Italian Legislative Decree n° 26 of 2014 (Authorization No. 12/2022-UT). The animals were used according to Directive 2010/63/EU regarding the protection of animals used for experimental or other scientific purposes, enforced by the Italian Legislative Decree n° 26 of 2014 and Ministerial Decree. Phase A authorization: 1017/2024-PR.

Expression and purification of MPXV OPG153 globular domain

FreeStyle 293-F cells (ThermoFisher) were cultured in FreeStyle 293 expression medium (Gibco) at 37 °C in a shaking incubator with 8% $CO_2$ and 80% humidity. For passaging, cells were seeded at a density of $0.3 \times 10^6$ cells/mL. Codon-optimized (GenScript) sequences for OPG153 (residues 1–366) from MPXV-Ib (NCBI Reference Sequence: NP_536565.1), VACV (NCBI Reference Sequence: NC_006998.1), and CPXV (GenBank: CAD90694.1) were cloned into the mammalian expression vector pαH, upstream of a human rhinovirus 3C (HRV3C) protease cleavage site, an 8 × His tag, and a Twin-Strep tag. The OPG153 plasmids were transiently transfected into FreeStyle 293-F cells at a density of $1 \times 10^6$ cells/mL using 25 kDa linear polyethylenimine (PEI) at a 3:1 PEI:DNA ratio in Opti-MEM (Gibco), with a final concentration of 0.1% Pluronic Acid F-68 (MP Biomedicals). Cells were cultured for 5–6 days at 37 °C in a shaker incubator with 8% $CO_2$ and 80% humidity. Culture supernatant was collected by centrifugation and filtered (0.22 μm) before being purified using Strep-Tactin XT 4Flow resin, following the manufacturer's instructions (IBA). The elution fraction was concentrated to <500 μL and purified via a Superdex 200 Increase 10/300 GL column (Cytiva) equilibrated in PBS. Protein-containing fractions were concentrated, flash-frozen, and stored at –70 °C.

Peripheral blood mononuclear cell isolation and memory B cell sorting

PBMC isolation was performed as previously described (*38*). Briefly, PBMCs were collected from heparin-treated whole blood by density gradient centrifugation (Ficoll-Paque PREMIUM, GE Healthcare). Then, PBMCs were incubated at room temperature with Live/Dead Fixable Aqua (Invitrogen) diluted 1:500 in 1× phosphate-buffered saline (PBS, Gibco). After 20 min of incubation, cells were washed with PBS and nonspecific binding was saturated with 50 μL of 20% normal rabbit serum (Life Technologies) diluted in PBS. Following incubation at 4 °C for 30 min, cells were stained with CD19 BV421 (BD Biosciences), IgM PerCP-Cy5.5 (BD Biosciences), CD27 PE (BD Biosciences), IgD Alexa Fluor 700 (BD Biosciences), CD3 PE-Cy7 (BioLegend), CD14 PE-Cy7 (BioLegend), and

CD56 PE-Cy7 (BioLegend) at 4 °C for 30 min. To identify OPG153$^{+}$ memory B cells (MBCs), cells were also stained for 20 min at 4 °C with OPG153 (residues 1–366) labeled with Strep-Tactin®XT DY-488 (IBA-Lifesciences). Following additional washing with PBS, cells were resuspended in Sorting Buffer (PBS + 2.5 mM EDTA). Stained MBCs (CD3$^{-}$CD14$^{-}$CD56$^{-}$CD19$^{+}$CD27$^{+}$IgD$^{-}$IgM$^{-}$) were single-cell sorted with a BD FACSAria™ Fusion (BD Biosciences) into 384-well microplates containing 3T3-CD40L feeder cells. Then, sorted memory B cells were incubated for 10–14 days with IL-2 and IL-21, as previously described (*38, 44*).

Single cell RT-PCR, Ig gene amplification and cloning of variable region genes

The whole process for monoclonal antibody (mAb) heavy and light chain recovery and amplification was performed as previously described (*38, 39*). In brief, from the original 384-well sorting plate, 5 µL of cell lysate of selected antibodies was used to perform RT-PCR, with random hexamers primers, using Superscript IV reverse transcriptase (Invitrogen). Heavy (VH) and light (VL) chain amplicons were obtained via two rounds of PCR. PCR products were purified using MultiScreen-PCRµ96 filter plates (Millipore) and then ligated into the corresponding predigested vectors using Gibson Assembly (NEB). Ligation products were used to transform One Shot® TOP10 Chemically Competent *E. coli* (Invitrogen), following manufacturer's instructions. Selected colonies were analyzed by plasmid isolation/restriction and sequencing. Large-scale isolation of plasmid DNA from recombinant *E. coli* cultures was performed using NucleoBond® Xtra Midi/Maxi - High copy plasmid purification (MACHEREY-NAGEL). Plasmids were quantified by NanoDrop™ and stored at –20 °C prior to use.

Expression and purification of human monoclonal antibodies

Expi293™ Expression System was used to transiently transfect Expi293F™ cells with plasmids carrying the antibody heavy chain and light chain with a 1:2 ratio, according to the manufacturer's protocol (ThermoFisher). Following harvest and clarification by centrifugation, cell culture supernatants were filtered through a 0.22-µm filter. The antibodies were then purified by affinity chromatography using the ÄKTA go™ chromatography system (Cytiva) at room temperature. Specifically, filtrated culture supernatants were passed over a 1–5 mL HiTrap Protein G HP column (Cytiva) previously equilibrated in Buffer A (0.02 M $NaH_2PO_4$, pH 7). Monoclonal antibodies were eluted by applying a step elution of 6 CV of Buffer B (0.1 M glycine-HCl, pH 2.7). Elution steps were collected in 1 mL fractions, which were analyzed by nonreducing SDS-PAGE. Fractions showing the presence of IgG were pooled and dialyzed into PBS overnight at 4 °C. Final antibody concentrations were determined by measuring the A520 using Pierce BCA Protein Assay Kit (Thermo Scientific). Purified proteins were stored at –80 °C prior to use.

Viral strains

The MPXV Clade IIb strain used to perform the cytopathic effect-based microneutralization assay (CPE-MN) neutralization assay was obtained from the European Virus Archive – GLOBAL (EVAg) (Monkeypox virus, 2022, Slovenia, ex Gran Canaria; Ref-SKU: 005V-04714). The VACV strain used to perform the U2OS viral-mediated cell-cell fusion assay was obtained from American Type Culture Collection (ATCC; Ref. ATCC-VR-1549 lotto 70047009). The CPXV strain used for viral propagation and ELISA was CPXV Taunton (UK) (*57*). The MPXV Clade Ib strain was isolated from a man in his mid-30s with no history of orthopoxvirus vaccination who had traveled between Sweden and Africa in August 2024 (*58, 59*). The isolate MpxV/PHAS-506/Passage- 03/SWE/2024_09_11, Clade Ib has been provided by the Public Health Agency of Sweden to improve the quality of diagnostics relevant for infectious disease control, treatment and/or other studies of relevance for public health.

Viral propagation

The MPXV Clade IIb, VACV, and CPXV viruses were propagated in Vero E6 cells (ATCC 1586) cultured in Dulbecco's Modified Eagle Medium (DMEM; Gibco) high glucose with 2% fetal bovine serum (FBS; Gibco), 100 U/mL penicillin (Gibco), and 100 mg/mL streptomycin (Gibco). Briefly, Vero E6 were seeded at a density of $1.8 \times 10^5$ cells/mL in T175 flasks and incubated at 37 °C, 5% $CO_2$ for 18–20 h. Then, cells were incubated with MPXV Clade IIb, VACV, or CPXV at a multiplicity of infection (MOI) of 0.001. After 48 h, cells and cell supernatant were scraped in 1 mL of culture medium, recovered, and subjected to three freeze-thaw cycles. Next, infected cells and supernatant were pelleted at 469 × *g* for 5 min at 4 °C and stored at –80 °C. For MV and EV MPXV isolation, the Vero E6 infection was performed as described above. Then, to isolate the EV MPXV form, cell supernatant was collected and stored at 4 °C. In parallel, to collect the MV MPVX form, Vero E6 infected cells were scraped in 10 mL of culture medium, subjected to three freeze-thaw cycles, and pelleted at 469 × *g* for 5 min. The resulting supernatant containing the MV form was stored at –80 °C.

MPXV Clade IIb MPXV, EV, and MV neutralization assays

Neutralization assays with authentic MPXV were performed in the biosafety level 3 (BSL3) laboratories at Toscana Life Sciences in Siena (Italy). BSL3 laboratories are approved by a Certified Biosafety Professional and are inspected annually by local authorities. To evaluate the neutralization activity and identify nAbs, sorting supernatants were screened in a CPE-MN assay (*38, 39*). In brief, pre-diluted (1:4) sorting supernatants were incubated with an MPXV viral solution containing 25 median tissue culture infectious dose ($25TCID_{50}$) of virus, in the presence of a final concentration of 3% baby rabbit complement (BRC). After a 1 h incubation at 37 °C, 5% $CO_2$, the mixture was added to a 96-well plate containing a sub-confluent Vero E6 cell monolayer. Plates were incubated for 4 days at 37 °C in a humidified environment with 5% $CO_2$, then examined for CPE by means of an inverted optical microscope by two independent operators. To evaluate the neutralization potency of purified mAbs, plasma, and serum samples, the assay was performed as described above but using a viral solution of $100TCID_{50}$ of MPXV or its MV or EV forms. EV or MV forms were incubated at 37 °C with 5% $CO_2$ in the presence of a final concentration of 3% BRC and with an anti-MV (MPXV-26) or anti-EV (VACV-22) nAb (*20*), respectively. All mAbs were tested at two-fold serial dilutions starting at 200 µg/mL, while human and mouse plasma samples were heat-inactivated for 30 min at 56 °C and tested at two-fold serial dilutions starting at a 1:10 dilution.

Enzyme-linked immunosorbent assay (ELISA)

ELISAs utilizing MPXV, VACV, or CPXV were all performed using the same method, with variation in the coating virus. Briefly, ELISA plates (Greiner) were coated with UV-inactivated (0.500 mJ/cm$^2$) virus diluted 1:10 in PBS and incubated at 4 °C overnight. Then, plates were washed three times with PBS-0.05% Tween-20 before blocking at 37 °C with 50 µL/well of saturation buffer (PBS + 1% BSA). After washing (PBS supplemented with 0.05% Tween-20), plates were incubated for 1 h at 37 °C with mAbs (10 µg/mL) diluted in dilution buffer (PBS + 1% BSA + 0.05% Tween-20). After washing as above, 25 µL/well of alkaline-phosphatase-conjugated Goat Anti-Human IgG (Southern Biotech:Cat# 2040-01) diluted 1:2,000 in dilution buffer was added and plates were incubated at 37 °C for 1 h. After washing, plates were incubated with 25 µL of p-nitrophenyl phosphate (PnPP; Sigma) and the reaction was measured at a wavelength of 405 nm using the Varioskan Lux Reader (Thermo Fisher Scientific). An unrelated mAb was used as a negative control and sample buffer was used as a blank. The threshold for sample positivity was set at 2-fold the blank OD. Technical duplicates were performed, and all data were analyzed using GraphPad Prism Software (version 8.4.2).

MPXV Clade Ib neutralization assay

The neutralization assay was conducted as previously described (*40*). Briefly, U2OS cells were plated at $8 \times 10^3$ cells/well in µClear 96-well plates (Greiner Bio-One). The following day, in a BSL3 facility, each virus was incubated with 1% of human serum as a source of complement (pool of two non-neutralizing sera) and serial antibody dilutions, starting at 25 µg/mL. After two hours, the complement mix was added to the cells. The viral inoculum was determined to obtain a non-saturating infection (*40, 58*). Forty-eight hours later, cells were fixed for 30 min at room temperature with 4% paraformaldehyde (PFA, Electron Microscopy Sciences), washed and immunostained for MPXV antigens with rabbit polyclonal anti-VACV antibodies (PA1-7258, Invitrogen), and an Alexa Fluor 488-coupled Goat anti-Rabbit antibody (Invitrogen). Nuclei were stained with Hoechst (1:10,000; Invitrogen). Images were acquired with an Opera Phenix high-content confocal microscope (PerkinElmer). For each condition, infection was quantified by calculating the total area of MPXV-positive cells ($MPXV^+$ area) and the nuclei were counted using the Harmony software (PerkinElmer). The percentage of infection inhibition was calculated from the $MPXV^+$ area using the following formula:

$$100 * \left[1 - \left[\frac{(MPXV^{+}\ area\ with\ Antibody) - (mean\ area\ of\ 'non - infected'controls)}{(mean\ area\ of\ 'no\ antibody'\ infected\ controls) - (mean\ area\ of\ 'non - infected'controls)}\right]\right]$$

Inhibition activity of each antibody was expressed as the $IC_{50}$ (half-maximal inhibitory concentration). $IC_{50}$s were calculated based on an inhibitory dose-response curve with a variable slope model, using the percentage of inhibition at the different antibody concentrations. An unrelated mAb was used as a negative control.

U2OS VACV-mediated cell-cell fusion assay

VACV neutralization assays were performed in U2OS-GFP 1–10 and U2OS GFP 11 cells carrying a GFP-Split complementation system (*41*). U2OS cells were grown in DMEM supplemented with 10% FBS, 100 U/mL penicillin, 100 mg/mL streptomycin, 1 µg/mL puromycin, and 10 µg/mL blasticidin. U2OS-GFP 1–10 and U2OS GFP 11 cells were mixed (1:1) and $8 \times 10^3$ cells/well were plated in a 96-well plate (µClear) 24 h before infection with authentic virus. Then, antibodies (serially diluted 1:2) were incubated with VACV viral solution containing the appropriate MOI in the presence of 2.5% of BRC in a final volume of 100 µL. The viral inoculum was determined to obtain a non-saturating infection (*40*). After a 1 h incubation at 37 °C, 5% $CO_2$, the antibody-virus mix was added to the U2OS cells. After a 72 h incubation at 37 °C in a humidified environment, cells were fixed in 4% PFA (Scientific Chemicals) for 30 min at room temperature, washed, and stained for 20 min with DAPI 62248 (1:2,000). Images were acquired on an Opera Phenix High Content Screening System (PerkinElmer) and analyzed with Harmony High-Content Imaging and Analysis Software (version 4.9). All mAbs were tested at 200 µg/mL, while human and mouse plasma samples were heat-inactivated for 30 min at 56 °C and tested at two-fold serial dilutions starting at 1:10 dilution. The threshold of sera neutralization was set at 50% the reduction of syncytia formation. The neutralization potency of each mAb was calculated as the percentage of syncytia reduction compared to the control of infection. Then the $IC_{50}$ was calculated with GraphPad Prism Software (version 8.4.2). Technical triplicates were performed.

Antibody-dependent cellular phagocytosis (ADCP) assay with inactivated MPXV

A flow cytometry-based assay was used to analyze antibody-dependent cellular phagocytosis (ADCP). Briefly, UV-inactivated MPXV was labelled using Alexa Fluor™ 488 NHS Ester (Succinimidyl Ester) kit and coated with the Dynabeads™ Intact Virus Enrichment (Catalog Numbers 10700D, Invitrogen), following the manufacturer's instructions. The virus-beads were incubated with nAbs or heated-inactivated plasma diluted in complete RPMI (Gibco) (1:40) for 1 h at room temperature. This mix was incubated with the monocytic THP-1 cell line for 6 h at 37 °C, 5% $CO_2$. Next, cells were fixed with fixation buffer (BD Biosciences) and acquired on BD CANTO (Becton Dickinson). Single technical replicates were performed for each experiment. Unrelated plasma and an unrelated mAb were used as negative controls. The phagocytosis score was calculated as the percentage of THP-1 cells that engulfed fluorescent beads multiplied by the median fluorescence intensity of the population. The threshold for sample positivity was set at 2-fold the negative control. Data were collected with the BD FACSDiva Software v9.0 (BD Biosciences) and analyzed with FlowJo™ Software (version 10).

Antibody-dependent cellular cytotoxicity (ADCC)

A CD16 activation reporter assay was performed to analyze ADCC activity (*60*). High-binding 96-well plates were coated with UV-inactivated MPXV diluted in PBS and incubated overnight at 4 °C. After incubation, the plate was washed 3 × with PBS supplemented with 0.01% Tween-20 and blocked with PBS supplemented with 2.5% BSA for 1 h at room temperature. Then, plasma or mAbs were diluted in growth medium (IMDM, 2 mM L-glutamine, 25 mM HEPES, 10% FBS) and added to the washed plate. After 30 min at room temperature, 100,000 Jurkat Lucia NFAT CD16 cells/well (Invivogen) were added. Antibodies and cells were incubated for 24 h at 37 °C with 5% $CO_2$. After incubation, 25 µL of cell supernatant was transferred into 96-well white walled clear bottom polystyrene plates (Costar) and mixed with 75 µL of reconstituted QUANTI-Luc reagent (InvivoGen). Luminescence was immediately read using a Varioskan plate reader (Molecular Devices). The assay was performed with two biological replicates. The threshold for sample positivity was set at 4-fold the negative control. Unrelated plasma and an unrelated mAb were used as negative controls.

AlphaFold 3-based predictive modeling

Amino-acid sequences for 40 full-length MPXV envelope proteins and the antibody variable domains of 12 mAbs were prepared in FASTA format. Three entities were defined: (1) full-length antigen, (2) VH, and (3) VL (stoichiometry 1:1:1; entity types "protein"). Model predictions were run on the online AlphaFold 3 Server (*37*) (accessed October 2024) in protein–protein complex mode with, yielding 5 predictions each for a total of 480 predictions. All jobs used default AlphaFold 3 databases and presets, with explicit random seeds recorded per run. Predictions were ranked by the interface predicted template modeling (ipTM) score provided by AlphaFold 3, defining an ipTM >0.80 as a high-confidence prediction. Analysis in ChimeraX (*71*) was used to assess the plausibility of the predicted VH/VL–antibody (i.e. the absence of steric clashes, and the presence of hydrogen bonds, hydrophobic contacts, electrostatic interactions, and/or van der Waals forces). Per-residue confidence (predicted Local Distance Difference Test, pLDDT) was also used to assess the confidence of the VH/VL–antigen interface.

Commercial expression of VARV OPG153 globular domain

Residues 1–366 of VARV OPG153 (NCBI Reference Sequence: NP_042177.1) with a C-terminal 8 × His tag were provided to GenScript for plasmid synthesis, protein expression, and affinity purification. Ni-NTA elution was buffer exchanged into PBS for shipping. The received protein preparation was concentrated to <500 µL and purified using a Superdex 200 Increase 10/300 GL column (Cytiva) equilibrated in PBS. Protein-containing fractions were concentrated, flash frozen, and stored at –70 °C.

OPG153 flow cytometry-based binding assay

OPG153 1–366 with a C-terminal 8 × His-tag from MPXV, VACV, or CPXV was used to coat magnetic beads (Dynabeads His-Tag, Invitrogen) according to the manufacturer's instructions. OPG153-beads were incubated with 20 µg/mL of nAbs diluted in PBS for 40 min at room temperature. To detect binding, beads were washed and stained with the Alexa Fluor 488-labeled Goat anti-Human IgG (H + L) secondary antibody (Invitrogen) diluted 1:2,000. After 30 min of incubation, beads were washed, resuspended in 150 µL of PBS, and analyzed using the BD CANTO (Becton Dickinson). Beads incubated with an unrelated mAb were used as a negative control. Data were collected with the BD FACSDiva Software v9.0 (BD Biosciences) and analyzed with FlowJo™ Software (version 10).

OPG153 flow cytometry-based competition assay

MPXV OPG153 1–366 with a C-terminal 8 × His-tag was used to coat magnetic beads (Dynabeads His-Tag, Invitrogen) according to the manufacturer's instructions. OPG153-beads were incubated with 20 µg/mL of nAbs diluted in PBS for 40 min at room temperature. The mixture was washed with 100 µL of PBS and incubated with labeled 12I12 or 08E11 antibodies. Following a 40-minute incubation at room temperature, the mixture was washed with PBS, resuspended in 150 µL of PBS-1% BSA, and analyzed using the BD CANTO (Becton Dickinson). Beads incubated with unlabeled plus labelled 12I12 or 08E11 nAbs were used as a positive control. Beads without OPG153 antigen or beads incubated with an unrelated mAb were used as negative controls. FACSDiva Software (version 9) was used for data acquisition. Analysis was performed using FlowJo (version 10).

Functional repertoire analyses

The VH and VL sequence reads of nAbs were manually curated and retrieved using CLC Sequence Viewer (Qiagen). Aberrant sequences were removed from the dataset. Analyzed reads were saved in FASTA format and the repertoire analyses were performed using Cloanalyst (http://www.bu.edu/computationalimmunology/research/software/) (*61*).

Antibody expression and purification for structural studies

Plasmids encoding antibody genes were transiently co-transfected at a 1:1 heavy-to-light chain ratio into FreeStyle 293-F cells, following the same protocol used for OPG153 globular domain expression. Culture supernatant was collected by centrifugation, filtered (0.22 µm), and purified using a column packed with Pierce Protein A Plus Agarose resin (ThermoFisher) equilibrated in PBS. The resin was washed with PBS, and antibodies were eluted with 100 mM glycine-HCl (pH 3.0), then immediately neutralized with 1:10 volume of 1 M Tris-HCl (pH 8.0). Eluted antibodies were buffer-exchanged into PBS, concentrated, flash-frozen, and stored at –70 °C.

Fab preparation

Purified IgG was digested with LysC (Fisher Scientific) at a 1:1,000 molar ratio (IgG:LysC) overnight at 37 °C followed by quenching with cOmplete Protease Inhibitor Cocktail (Sigma-Aldrich). The digested IgG was passed over CaptureSelect IgG-CH1 affinity resin (ThermoFisher), washed with PBS, eluted with 100 mM glycine-HCl (pH 3.0), then immediately neutralized with 1:10 volume of 1 M Tris-HCl (pH 8.0).

Surface plasmon resonance

An anti-StreptagII IgG antibody (*62*) was immobilized on a CM5 sensor chip (Cytiva, cat #BR100012) using amine coupling chemistry following the standard Biacore X100 (GE Healthcare) manufacturer's protocol. Immobilization was performed for 7 min, reaching ~5000 RU. A ligand (twin-Strep-tagged OPG153 globular domain protein) was subsequently captured on flow cell 2 to ~300 RU. All analytes (Fab) were titrated in a two-fold dilution series from 200 nM to 1.5625 nM, with the exception of the 12I12/VACV OPG153 run, which ranged from 800 nM to 6.25 nM. Each dilution was flowed over both flow cells 1 and 2. The sensor surface was regenerated using 0.1 % SDS and 10 mM glycine (pH 2.0) between cycles. Binding curves were double-reference-subtracted and fitted to a 1:1 binding model using Biacore X100 software and plotted in GraphPad Prism (version 10.3.1).

Cryo-EM sample preparation

The MPXV OPG153 globular domain protein was diluted to a final concentration of 0.28 mg/mL (OPG153-08E11-12I12) or 0.22 mg/mL (OPG153-02M12) and pre-incubated with a 1.2-fold molar excess of 08E11 Fab and an equimolar amount of 12I12 Fab (OPG153-08E11-12I12) or an equimolar amount of 02M12 (OPG153-02M12) in 20 mM Tris (pH 8.0), 200 mM NaCl for 30 min at room temperature. Amphipol A8-35 (Anatrace, cat. #A835) was added for a final concentration of 0.02% immediately before freezing. A 3.8 µL aliquot of the mixture was applied to a 30-s glow-discharged (PELCO easiGlow™) UltrAuFoil 1.2/1.3 300-mesh TEM grid (Electron Microscopy Sciences, cat. #Q350AR13A). The grid was plunge-frozen in liquid ethane using a Vitrobot Mark IV (FEI) with a blot time of 9 s, blot force of 1, at 22 °C and 100% humidity.

Cryo-EM data collection, processing, and analysis

The TEM grids were loaded onto a 200 kV Glacios (Thermo Fisher Scientific) equipped with a Falcon 4 detector. All movies were collected at a total dose of 50 $e^{-}/Å^{2}$ under 150,000× magnification corresponding to a pixel size of 0.933 Å/pix with a defocus range of –1.5 to –2.5 µm. Data collection was automated using SerialEM (*63*). A total of 890 movies at 0° and 1,147 movies at 30° were collected for the OPG153-08E11-12I12 dataset, and 299 movies at 0° and 1,444 movies at 30° were collected for the OPG153-02M12 dataset. Datasets were imported into CryoSPARC v4.5.3 (*64*) for processing. The final maps for model building were orientation-rebalanced and then sharpened using DeepEMhancer (*65*). Detailed EM data processing workflows are shown in **figs. S6** and **7**. The AlphaFold 3-predicted models were used as a starting point for model building. Iterative refinement was performed using Phenix 1.21.2-5419 real-space refinement (*66, 67*), Coot 0.9.8.95 EL (CCP4) (*68*), and UCSF ChimeraX-1.9 ISOLDE (*69*). Total buried solvent accessible surface area was calculated with PDBePISA (*70*). Detailed cryo-EM data collection and model statistics are provided in **table S3**.

Production of hyperimmune serum in immunized mice

Three groups of ten female BALB/c mice (7–8 weeks of age at study initiation) were immunized by intramuscular (IM) injection on days 0 and 21. Mice in two of the groups were immunized with 10 µg of recombinant protein—either OPG153 (residues 1–366) or SARS-CoV-2 S protein HexaPro (*55*)—formulated in 25 µL of AddaVax™ (Squalene-based oil-in-water adjuvant, InvivoGen) and diluted in PBS to a final volume of 50 µL. Mice in a third group received 50 µL of MVA-BN vaccine. On day 35, terminal blood samples were collected via exsanguination under isofluorane anesthesia for serum processing. Sacrifice was carried out by cervical dislocation. Targeted volume for terminal blood collection was approximately 800 µL per mouse. Serum was isolated from the whole blood by centrifugation and stored in vials at –20 °C until usage.

Figure preparation

Molecular graphics and analyses were performed using UCSF ChimeraX (*71*). Cryo-EM map values were normalized for figure preparation to mean = 0 and $\sigma = 1$ in UCSF ChimeraX using the 'volume scale' function. Cryo-EM maps are contoured at $6\sigma$ in all images, unless otherwise noted. R.M.S.D. was measured in UCSF ChimeraX using the Matchmaker tool. EM maps were colored using the Color Zone tool in UCSF ChimeraX with a 3–5 Å radius. Figures were compiled in Adobe Illustrator (Adobe). Cartoon illustrations (human silhouette, mouse, and syringe) were made using generative AI in Adobe Illustrator (Adobe).

Statistical analysis

Statistical analysis was performed using GraphPad Prism Software (version 8.4.2). Statistical significance is shown as *, **, *** and **** for p values ≤ 0.05, ≤ 0.01, ≤ 0.001 and ≤ 0.0001 respectively. Significant differences between groups were evaluated using Tukey's multiple comparisons test one-way ANOVA (95% confidence interval). Serum neutralization titers are shown as $Log_{10}$. Black lines denote the median values for each group. Dotted lines represent the lower limit of detection (LLD).

**A**

**MPXV infection**

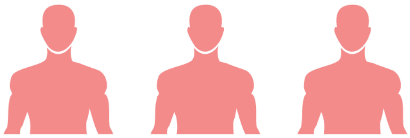

| Subject ID | Gender | Age (years) | MPXV infection (dd/month/yyyy) | Blood collection (dd/month/yyyy) | Days from vaccination to blood collection |
|---|---|---|---|---|---|
| INF-01S | Male | 47 | 09 July 2022 | 19 September 2022 | 72 |
| INF-02M | Male | 50 | 04 July 2022 | 20 January 2023 | 200 |
| INF-01V | Male | 44 | 02 November 2022 | 06 December 2022 | 33 |

**B**

**MVA-BN vaccination**

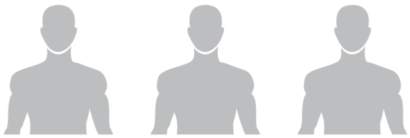

| Subject ID | Gender | Age (years) | MVA-BN vaccination (dd/month/yyyy) | Blood collection (dd/month/yyyy) | Days from vaccination to blood collection |
|---|---|---|---|---|---|
| VAC-01S | Male | 51 | 23 September 2022 | 30 January 2023 | 129 |
| VAC-03S | Male | 27 | 21 September 2022 | 09 February 2023 | 141 |
| VAC-04S | Male | 53 | 26 January 2023 | 17 April 2023 | 81 |

**C**

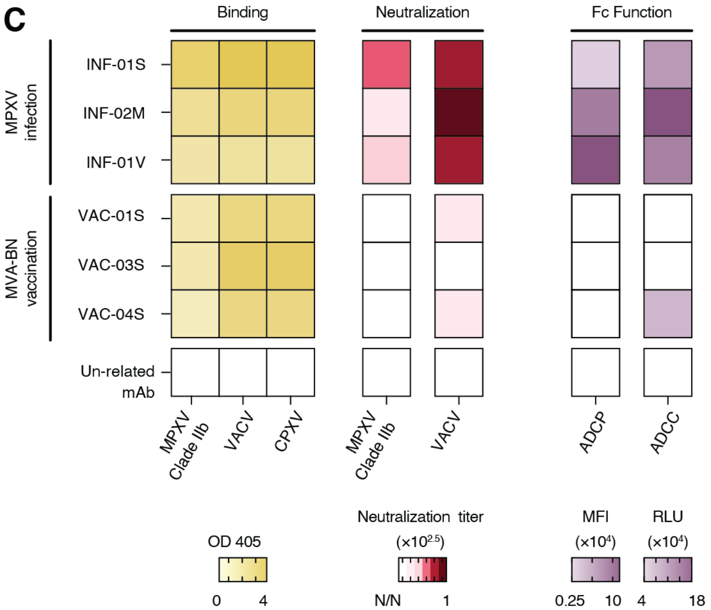


**Figure S1. Donor cohorts and characterization of the polyclonal antibody responses.** (**A**, **B**) Schematic overview of donor enrollment and sample collection for (A) MPXV-infected individuals (n = 3) and (B) MVA-BN vaccine recipients (n = 3). (**C**) Heatmaps summarizing polyclonal antibody responses elicited by infection or vaccination through binding assay with UV-inactivated orthopoxviruses (*left*), neutralization of MPXV and VACV (*middle*), and Fc-mediated effector functions (ADCP, ADCC) against UV-inactivated MPXV (*right*). An unrelated mAb was used as a control.

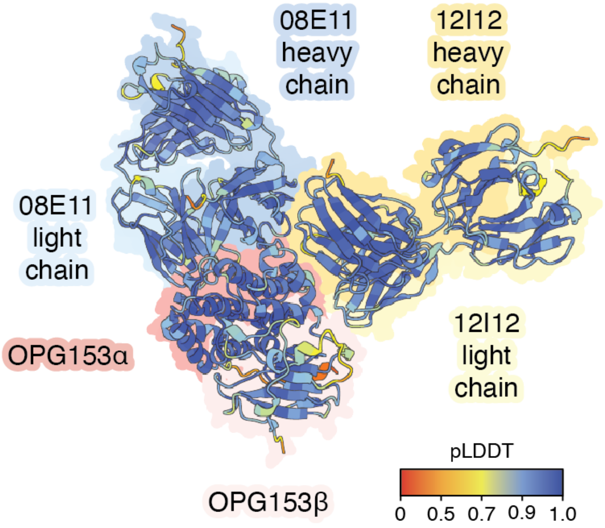


**Figure. S2. AlphaFold 3 modeling predicts non-overlapping binding of 08E11 and 12I12 to MPXV OPG153.** AlphaFold 3-predicted binding of the 08E11 and 12I12 Fabs to distinct, non-overlapping epitopes on MPXV OPG153, colored by predicted local distance difference test (pLDDT) scores. MPXV OPG153 residues 1–15 and 367–530 were omitted for clarity.

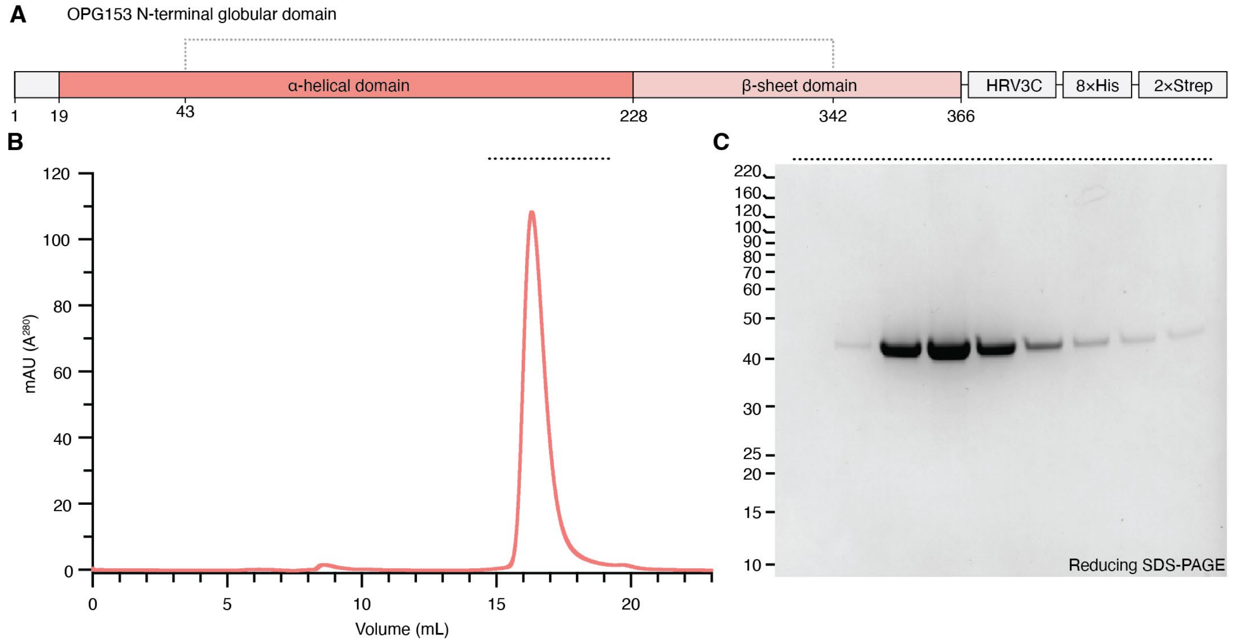


**Figure S3. Purification of the N-terminal globular domain of OPG153.** (**A**) Schematic representation of the OPG153 globular domain construct used for recombinant expression. The dotted line indicates a disulfide bond. (**B**) Recombinant MPXV OPG153 globular domain was purified by Strep-Tactin XT affinity chromatography, followed by Size-exclusion chromatography on a Superdex200 Increase 10/300 column equilibrated in PBS. Fractions corresponding to the dotted line were analyzed in (C). (**C**) Coomassie blue-stained reducing NuPAGE 4–12% Bis-Tris gel. Positions of the molecular weight standards are shown on the left, in kDa.

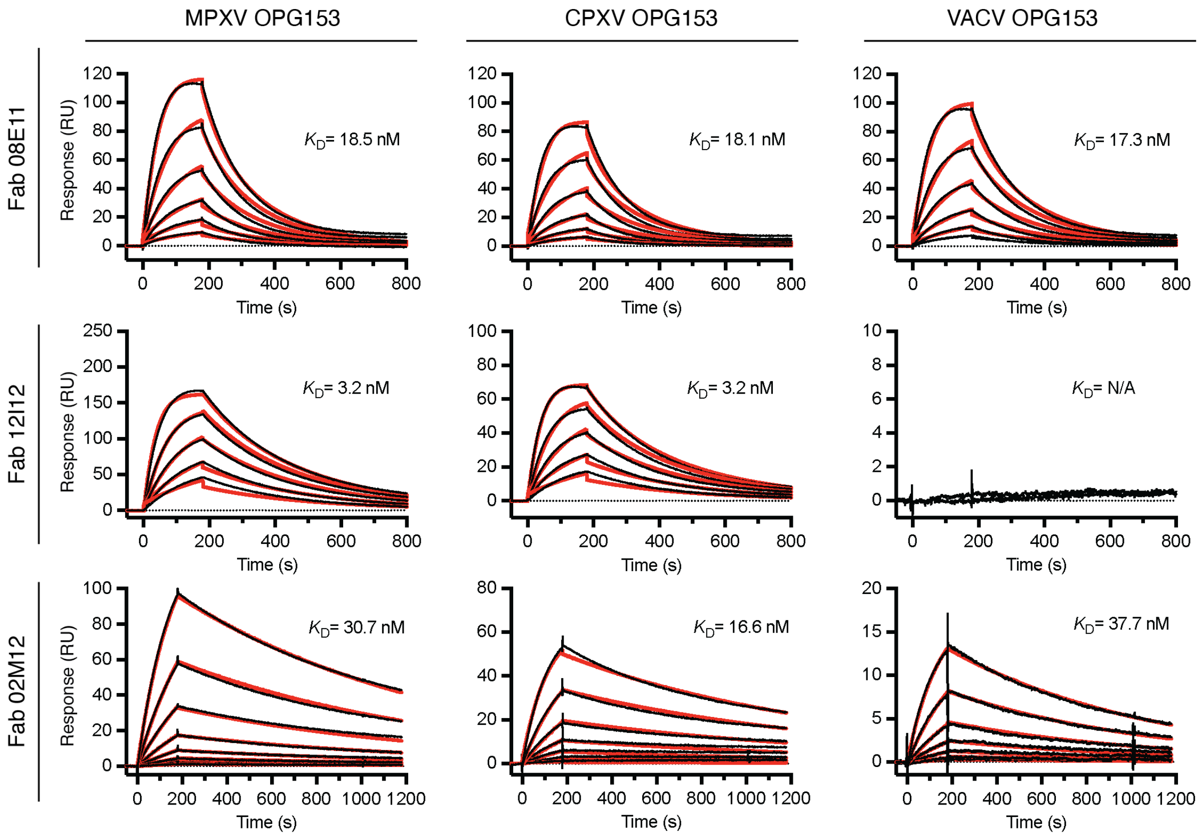


**Figure S4. Binding kinetics of 08E11, 12I12, and 02M12 Fabs to orthopoxvirus OPG153.** Surface plasmon resonance binding analysis of Fabs 08E11 (*top*), 12I12 (*middle*), and 02M12 (*bottom*) binding to the globular domains of OPG153 from MPXV (*left*), CPXV (*middle*), and VACV (*right*). Kinetic parameters were calculated to assess binding affinities. Experimental data (black) were fitted to a 1:1 binding model (red).

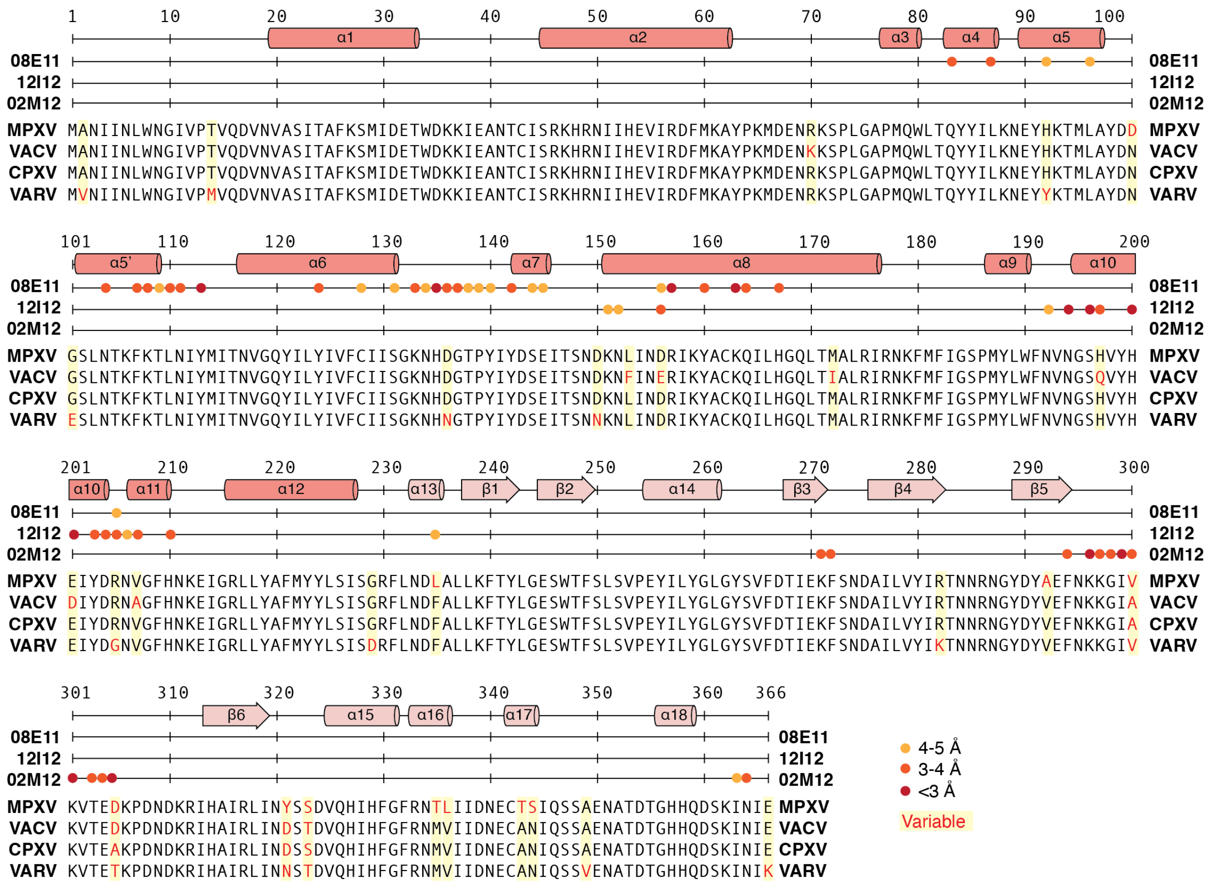

**Figure S5. Secondary structure and sequence alignment of orthopoxvirus OPG153.** Multiple sequence alignment of residues 1–366 of OPG153 from MPXV, VACV, CPXV, and VARV, with residues involved in antibody interactions indicated by colored circles. The secondary structure of OPG153 is annotated above the sequence alignment (*54*). Variable positions across orthopoxvirus sequences are highlighted in yellow.

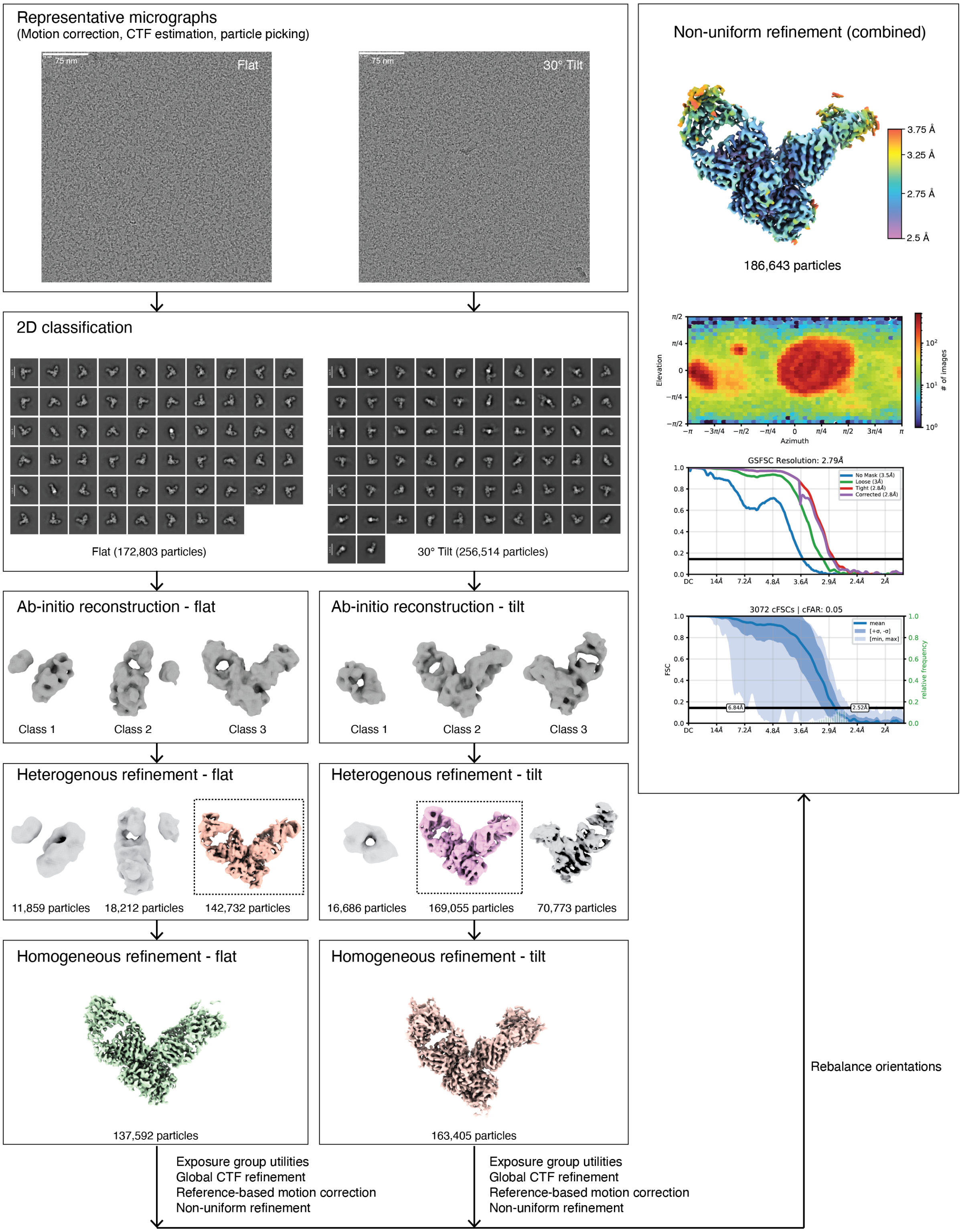


**Figure S6. Cryo-EM data processing workflow for the OPG153-08E11-12I12 complex.** Single-particle cryo-EM data were processed using CryoSPARC v4.5.3 (*64*). The workflow included motion correction, contrast transfer function (CTF) estimation, particle picking, 2D classification, *ab-initio* reconstruction, heterogenous and homogeneous refinement, reference-based motion correction, and non-uniform refinement. The final map was sharpened using DeepEMhancer (*65*).

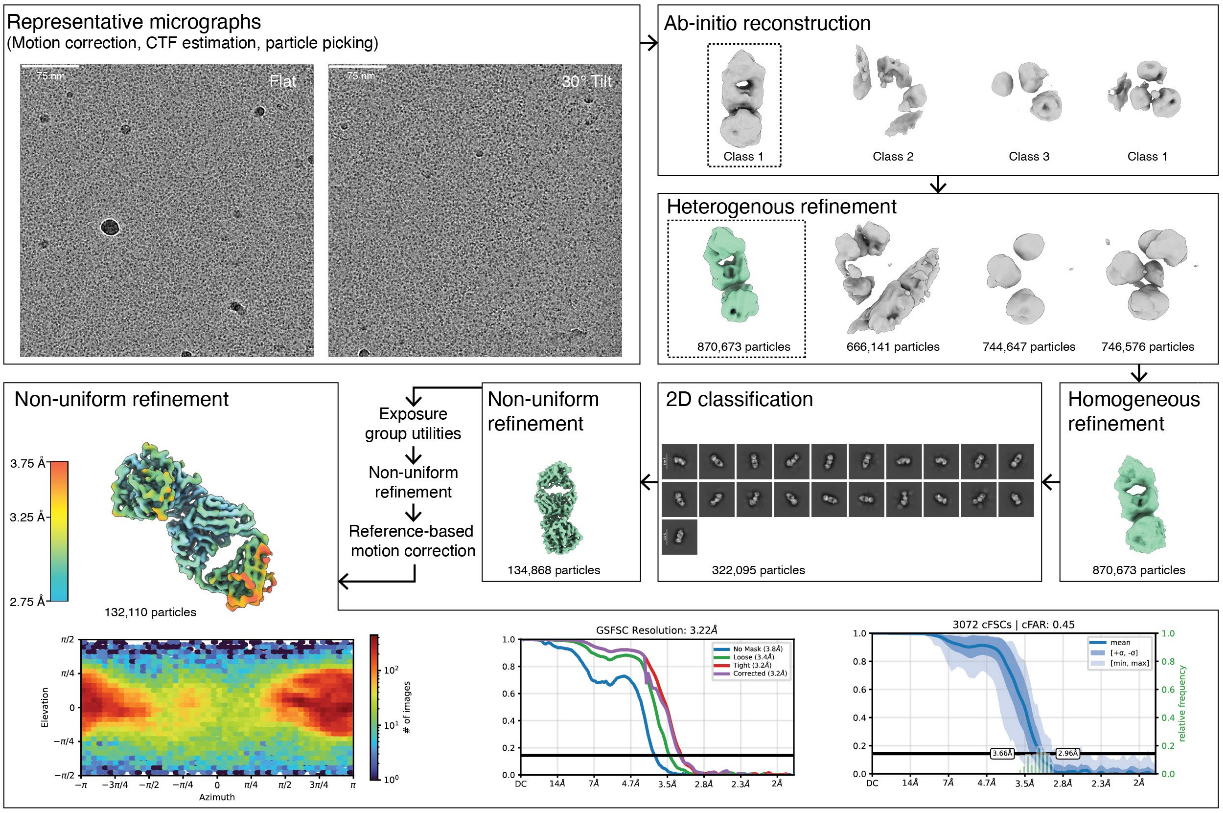


**Figure S7. Cryo-EM data processing workflow for the OPG153-02M12 complex.** Single-particle cryo-EM data were processed using CryoSPARC v4.5.3 (*64*). The workflow included motion correction, contrast transfer function (CTF) estimation, particle picking, 2D classification, *ab-initio* reconstruction, heterogenous and homogeneous refinement, reference-based motion correction, and non-uniform refinement. The final map was sharpened using DeepEMhancer (*65*).

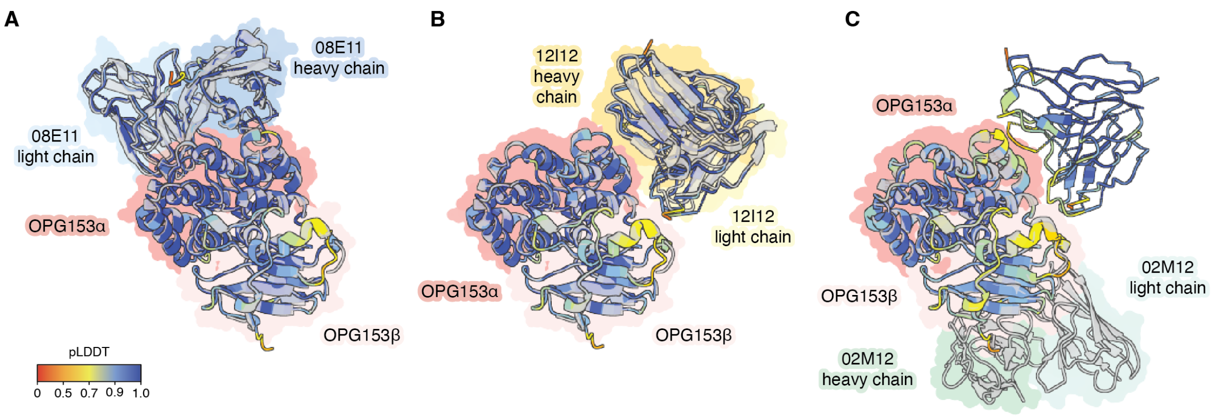


**Figure S8. Structural comparison of cryo-EM and AlphaFold 3 models.** (**A**–**C**) Overlays of cryo-EM models (transparent gray) and AlphaFold 3-predicted structures (colored by pLDDT scores) illustrate conformational agreement for the (A) OPG153-08E11 and (B) OPG153-12I12 complexes, but notable disagreement for (C) the 02M12 binding site on OPG153.

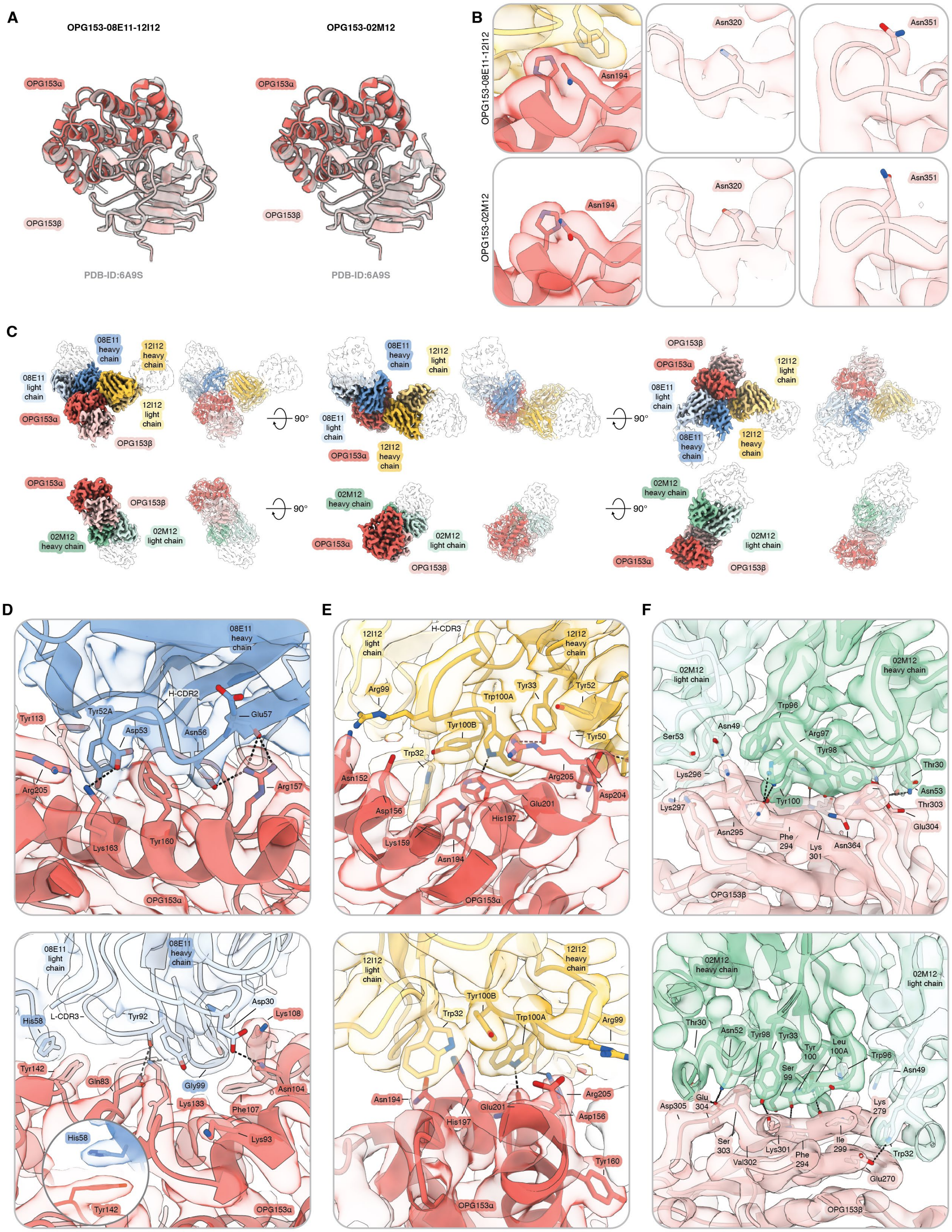


**Figure S9. Structural validation of OPG153-antibody complexes.** (**A**) Overlay of the cryo-EM structures of the MPXV OPG153 globular domain with the crystal structure of VACV OPG153 (PDB: 6A9S) (*54*). (**B**) Close-up views of cryo-EM maps at predicted N-linked glycosylation sites, contoured at 2σ. (**C**) Cryo-EM reconstructions of OPG153 in complex with 08E11 and 12I12 (*top*) or 02M12 (*bottom*), with domains colored as in **Fig. 2**. (**D**–**F**) Map-to-model agreement at the interfaces of MPXV OPG153 and nAbs (D) 08E11, (E) 12I12, and (F) 02M12, contoured at 6σ.

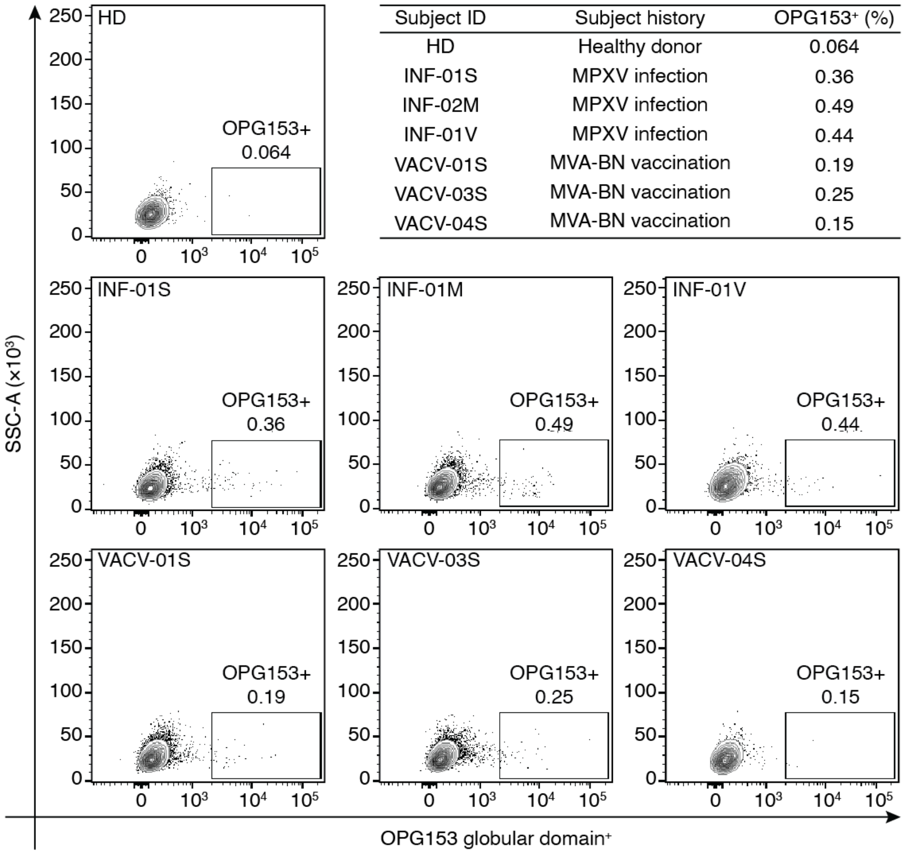


| Subject ID | Subject history | OPG153⁺ (%) |
|---|---|---|
| HD | Healthy donor | 0.064 |
| INF-01S | MPXV infection | 0.36 |
| INF-02M | MPXV infection | 0.49 |
| INF-01V | MPXV infection | 0.44 |
| VACV-01S | MVA-BN vaccination | 0.19 |
| VACV-03S | MVA-BN vaccination | 0.25 |
| VACV-04S | MVA-BN vaccination | 0.15 |

**Figure S10. Frequency of OPG153$^+$ MBCs elicited by MPXV infection or MVA-BN vaccination.** Flow cytometry dot plots showing the percentage of OPG153$^+$ class-switched memory B cells (CD19$^+$CD27$^+$IgD$^-$IgM$^-$) in each donor. Summary table displaying the frequencies of OPG153$^+$ MBCs in MPXV-infected individuals and MVA-BN vaccinees. PBMCs derived from a healthy donor (HD) were used as a negative control (*top right*).

**Table S1. Genetic characteristics of identified MPXV nAbs.**

| nAb ID | Clone ID | VH-JH usage | VH Mutation Freq (%) | HCDR3 Length (aa) | VL-JL usage | VL Mutation Freq (%) | LCDR3 Length (aa) |
|---|---|---|---|---|---|---|---|
| **01G09** | 1 | *IGHV3-33; IGHJ1-1* | 25.50 | 15 | *IGLV2-23; IGLJ3-1* | 27.50 | 13 |
| **02M12** | 2 | *IGHV1-2; IGHJ2-1* | 16.44 | 14 | *IGKV1-12; IGKJ2-1* | 20.63 | 9 |
| **02N10** | 3 | *IGHV4-39; IGHJ5-1* | 4.65 | 19 | *IGKV3-20; IGKJ5-1* | 0.72 | 5 |
| **04B17** | 4 | *IGHV3-9; IGHJ4-1* | 19.66 | 14 | *IGKV3-20; IGKJ1-1* | 23.49 | 6 |
| **04M05** | 5 | *IGHV3-30; IGHJ6-1* | 0.67 | 17 | *IGKV1-9; IGKJ2-1* | 1.84 | 9 |
| **08E11** | 6 | *IGHV3-30; IGHJ4-1* | 2.68 | 14 | *IGKV1-39; IGKJ2-1* | 23.83 | 9 |
| **08H03** | 7 | *IGHV3-23; IGHJ5-1* | 3.03 | 13 | *IGKV1-9; IGKJ2-1* | 1.84 | 9 |
| **09O02** | 8 | *IGHV1-2; IGHJ2-1* | 17.45 | 15 | *IGKV3-20; IGKJ4-1* | 23.83 | 8 |
| **12I12** | 9 | *IGHV4-30.4; IGHJ4-1* | 22.26 | 12 | *IGKV1-12; IGKJ1-1* | 24.91 | 9 |
| **13B11** | 10 | *IGHV2-5; IGHJ3-1* | 2.33 | 18 | *IGKV1-39; IGKJ4-1* | 4.04 | 9 |
| **13D15** | 11 | *IGHV3-7; IGHJ2-1* | 20.60 | 11 | *IGKV3-20; IGKJ1-1* | 3.61 | 8 |
| **16C15** | 12 | *IGHV3-43D; IGHJ4-1* | 19.66 | 15 | *IGLV2-14; IGLJ2-1* | 28.21 | 11 |

**Table S2. Neutralization potencies of identified nAbs.**

| mAb ID | MPXV-IIb ($IC_{100}$) - ng/mL | MPXV-Ib ($IC_{50}$) - ng/mL | MPXV-IIb MV ($IC_{100}$) - ng/mL | MPXV-IIb EV ($IC_{100}$) - ng/mL | VACV ($IC_{50}$) - ng/mL |
|---|---|---|---|---|---|
| **01G09** | 2,210.0 | 23,800.0 | N/N | N/N | N/N |
| **02M12** | 7,071.0 | 1,135.0 | 8,838.8 | N/N | N/N |
| **02N10** | 31.3 | 0.7 | 86.7 | 1,325.8 | 6.6 |
| **04B17** | 55.2 | 4.0 | 133.3 | 1,875.0 | 56.0 |
| **04M05** | 1,414.2 | 292.1 | 2,080.3 | 14,142.1 | 1,083.0 |
| **08E11** | 442.0 | 20.0 | 625.0 | 28,284.3 | 93.2 |
| **08H03** | 62.5 | 2.0 | 321.5 | 331.5 | 7.9 |
| **09O02** | 31.3 | 1.7 | 247.0 | 1,767.8 | 5.7 |
| **12I12** | 221.0 | 12.0 | 322.3 | N/N | N/N |
| **13B11** | 20,650.0 | 3,990.0 | N/N | N/N | N/N |
| **13D15** | 50,000.0 | 4,310.0 | 14,142.1 | N/N | N/N |
| **16C15** | 110.0 | 15.0 | 2,500.0 | 12,071.1 | 581.1 |
| **MPXV-26** | N/T | N/T | 9,451.3 | N/N | N/T |
| **VACV-22** | N/T | N/T | N/N | 40,000.0 | N/T |
| **Unrelated mAb** | N/N | N/N | N/N | N/N | N/N |

N/T= Not tested; N/N= Non-neutralizing

### Table S3. Cryo-EM data collection and refinement statistics.

| **EM data collection** | OPG153 - 08E11 Fab - 12I12 Fab | | OPG153 - 02M12 Fab | |
|---|---|---|---|---|
| Microscope | Glacios | | Glacios | |
| Voltage (kV) | 200 | | 200 | |
| Detector | Falcon 4 | | Falcon 4 | |
| Magnification (nominal) | 150,000 × | | 150,000 × | |
| Pixel size (Å/pix) | 0.93 | | 0.93 | |
| Exposure rate ($e^-$/pix/sec) | 2.24 | | 2.10 | |
| Exposure ($e^-$/Å$^2$) | 45 | | 49 | |
| Defocus range (µm) | 1.5–2.5 | | 1.5–2.5 | |
| Automation software | SerialEM | | SerialEM | |
| Sample | MPXV OPG153 at 0.28 mg/mL with 1.2 × molar excess Fab 08E11 and equimolar Fab 12I12 | | MPXV OPG153 at 0.22 mg/mL and equimolar Fab 02M12 | |
| Tilt angle (˚) | 0 | 30 | 0 | 30 |
| Micrographs collected | 890 | 1,147 | 299 | 1,444 |
| Micrographs used | 751 | 907 | 1,640 | |
| Particles extracted (total) | 653,530 | 1,004,306 | 3,028,037 | |

| **3D reconstruction statistics** | | |
|---|---|---|
| Particles | 186,643 | 132,110 |
| Symmetry | C1 | C1 |
| Map sharpening B-factor | –103.3 | –160.7 |
| Unmasked resolution at 0.5 FSC (Å) | 4.1 | 4.2 |
| Masked resolution at 0.5 FSC (Å) | 3.2 | 3.5 |
| Unmasked resolution at 0.143 FSC (Å) | 3.5 | 3.8 |
| Masked resolution at 0.143 FSC (Å) | 2.8 | 3.2 |

| **Model refinement and validation statistics** | | |
|---|---|---|
| Refinement package | ChimeraX ISOLDE, Phenix, Coot | Phenix, Coot |
| Refinement tool | Phenix real-space refinement | Phenix real-space refinement |
| Refinement strategies | min global, local_grid_search, adp, reference model restraints, ss restraints, rotamer restraints, Ramachandran restraints | min global, local_grid_search, adp, reference model restraints, ss restraints, rotamer restraints, Ramachandran restraints |
| Composition | | |
| Amino acids | 800 | 574 |
| RMSD bonds (Å) | 0.004 | 0.003 |
| RMSD angles (º) | 1.041 | 0.568 |
| Average B-factors | | |
| Amino acids | 59.1 | 66.6 |
| Ramachandran | | |
| Favored (%) | 97.7 | 98.2 |
| Allowed (%) | 2.3 | 1.8 |
| Outliers (%) | 0 | 0 |
| Rotamer outliers (%) | 0.86 | 0.80 |
| Clash score | 5.44 | 2.97 |
| C-beta outliers (%) | 0 | 0 |
| CaBLAM outliers (%) | 1.41 | 1.42 |
| CC (mask) | 0.79 | 0.79 |
| MolProbity score | 1.35 | 1.09 |
| EMRinger score | 3.60 | 3.57 |
